# Intermediate CA1 is Required for Object-in-Place Recognition Memory in Mice

**DOI:** 10.1101/2023.12.01.569628

**Authors:** Arely Cruz-Sanchez, Mehreen Inayat, Parjanya Parikh, Ryan Appings, Francesca Violi, Maithe Arruda-Carvalho

**Affiliations:** University of Toronto Scarborough, Department of Psychology, Toronto, Ontario, Canada, M1C1A4; University of Toronto, Department of Cell and Systems Biology, Toronto, Ontario, Canada, M1C1A4

## Abstract

Many behaviors that are essential for survival, such as retrieving food, finding shelter and locating predator cues, rely on forming effective associations between the identity and location of spatial elements. This identity-location association is commonly assessed in rodents using spontaneous object-in-place (OiP) recognition memory tasks. OiP recognition memory deficits are seen in autism spectrum disorder, schizophrenia, and are used to detect early onset of Alzheimer’s disease. These deficits are replicated in animal models of neurodevelopmental, neurodegenerative and chromosomal disorders. Mouse models have been widely adopted in behavioral and systems neuroscience research for their ease of genetic manipulations, and yet very few studies have successfully assessed OiP recognition memory or its neural correlates in mice. To address this limitation, we first established that adult C57/129J and C57BL/6J male and female mice are able to successfully perform the two-object, but not the four-object version of the spontaneous OiP recognition task, with retention intervals of five minutes and one hour. Next, using chemogenetic inhibition, we found that two-object OiP requires the activity of the intermediate CA1 (iCA1) subregion of the hippocampus, but not the medial prefrontal cortex or iCA1-medial prefrontal cortex connections. Our data identify hippocampal subregion specialization in the successful assessment of OiP recognition memory in mice, expanding our understanding of the neural basis of spatial memory processing.

**Significance Statement:** Associations between the identity and location of spatial elements (what-where associations), underlie essential behaviours such as finding food, locating shelter and safely navigating the environment. Deficits in identity-location processing occur in patients with neurodevelopmental and neurodegenerative disorders, and are replicated in rodent models using object-in-place (OiP) recognition tasks. While mice have emerged as a widely used animal model to study the biological mechanisms underlying these disorders, nothing is known about the neural substrates of OiP memory in mice. Here we have established and validated a robust experimental paradigm to assess OiP memory in mice, uncovering a specialized contribution of the hippocampal subregion intermediate CA1 to OiP performance and deepening our understanding of the neural signatures of spatial memory processing.

## Introduction

An animal’s ability to survive relies on the optimal navigation of their environment. This includes being able to *identify* and *locate* specific environmental cues, e.g. to find food and shelter or to avoid spaces previously visited by a predator. This ‘identity-location’ association is commonly tested in rodents through the object-in-place (OiP) recognition memory task (Poucet et al., 1986; Granon et al., 1996; Dix and Aggleton, 1999; Ameen-Ali et al., 2015). OiP recognition memory refers to the ability to recall an association between a previously encountered object and its specific location. Importantly, object-location association memory deficits are seen in people with schizophrenia (Wood et al., 2002; Burglen et al., 2004; Wout et al., 2006), and OiP memory deficits are reported in rodent models of neurodevelopmental and neurodegenerative disorders (Howland et al., 2012; Connors et al., 2014; Ballendine et al., 2015; Bonardi et al., 2016, 2021; Hall et al., 2016; Kentner et al., 2016).

In the most common version of the OiP task, rodents are first exposed to four distinct objects (Poucet, 1989; Dix and Aggleton, 1999; Ameen-Ali et al., 2015). Next, the position of two of those objects is switched (Figure 1A). OiP recognition memory is interpreted as increased exploration of the switched (or displaced) objects (Dix and Aggleton, 1999; Barker et al., 2007, 2017; Barker and Warburton, 2008, 2015; Ameen-Ali et al., 2015). An alternative version of this task exposes animals to two distinct objects, followed by the replacement of one of those objects with an identical copy of the remaining object (Eacott and Norman, 2004; Langston and Wood, 2010) (Figure 1G). In rats, the four-object version of the OiP task relies on the perirhinal cortex, hippocampus, medial prefrontal cortex (mPFC), mediodorsal thalamic nucleus, nucleus reuniens and some of their connections (Gaffan and Parker, 1996; Bussey et al., 2000; Good et al., 2007; Barker et al., 2007, 2020; Baxter et al., 2007; Bachevalier and Nemanic, 2008; Barker and Warburton, 2008; Griffiths et al., 2008; Barker and Warburton, 2009, 2011, 2015, 2018; Cross et al., 2013; Mitchnick et al., 2015, 2016, 2019; Savalli et al., 2015; Benn et al., 2016; Scott et al., 2017; Sabec et al., 2018), particularly between the hippocampal intermediate CA1 (iCA1) and the mPFC (Barker et al., 2017). While the two-object version of the OiP task in rats also relies on the hippocampus (Langston and Wood, 2010), hippocampal subregion and mPFC involvement in this version of the OiP task have not been investigated.

**Figure 1.**
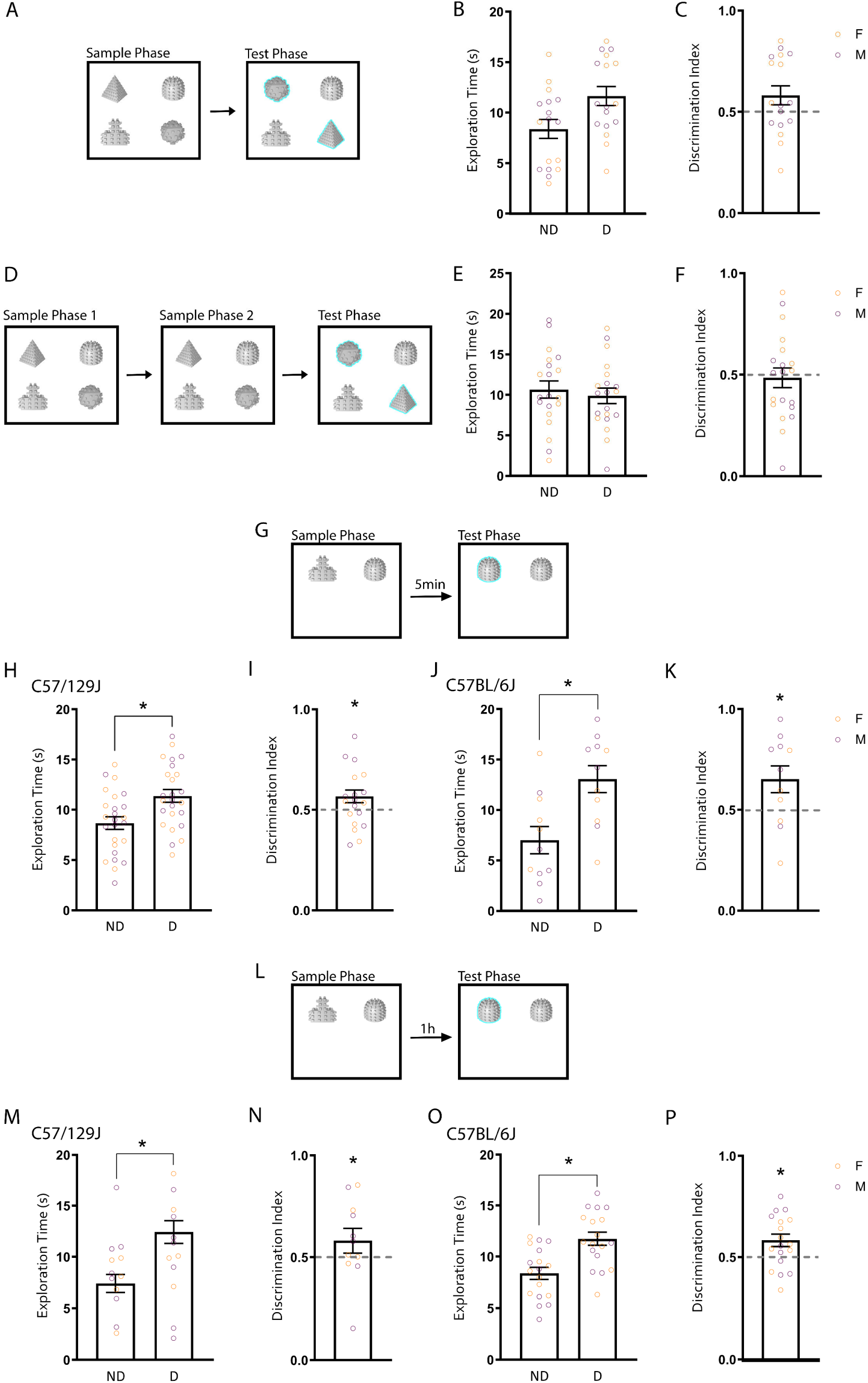
Mice show robust OiP memory across retention intervals in the two- but not four-object version of the OiP task. **A-C**. In the four-object, one-sample phase design (**A**), C57/129J mice underwent a sample phase in which they were exposed to four different objects. Five minutes later, they were exposed to the same objects in a test phase in which two of the objects switched locations (displaced objects; depicted with blue halo for representative purposes). During the test phase (**B**), there was no difference in exploration time between the displaced (D) and non-displaced (ND) objects. Calculation of a discrimination index for each animal (**C**) showed no difference from a chance level of preference (0.5). n= 17 (8 females, 9 males). **D-F**. In the four-object, two-sample phase design (**D**), C57/129J mice underwent two identical sample phases in which they were exposed to four different objects. Two minutes later, they were exposed to the same objects in a test phase in which two of the objects swapped locations (displaced objects; depicted with blue halo for representative purposes). During the test phase (**E**), the displaced (D) and non-displaced (ND) objects were explored equally. Calculation of a discrimination index for each animal (**F**) showed no difference from a chance level of preference (0.5). n= 20 (10 females, 10 males). **G-K**. In the two-object OiP task with a 5min retention interval (**G**), mice underwent a sample phase in which they were exposed to two distinct objects. Five minutes later, they were exposed to two objects in the same locations, but one of the objects was replaced by an identical copy of the remaining object (displaced object; depicted with blue halo for representative purposes). **H-I**. Two-object OiP data for C57/129J mice. During the test phase (**H**), C57/129J mice spent more time exploring the displaced (D) object compared to the non-displaced (ND) object. Calculation of a discrimination index for each animal (**I**) showed a significant preference for the displaced object over chance levels (0.5). n= 25 (12 females, 13 males). **J-K**. Two-object OiP data for C57BL/6J mice. During the test phase (**J**), C57BL/6J mice spent more time exploring the displaced (D) object compared to the non-displaced (ND) object. Calculation of a discrimination index for each animal (**K**) showed a significant preference for the displaced object over chance levels (0.5). n= 11 (5 females, 6 males). **L-P**. Two-object OiP task with a 1h retention interval. C57/129J (**M-N**) or C57BL/6J (**O-P**) mice underwent a sample phase in which they were exposed to two distinct objects. One hour later, they were exposed to two objects in the same locations, but one of the objects was replaced by an identical copy of the remaining object. C57/129J mice spent more time exploring the displaced (D) object compared to the non-displaced (ND) object (**M**). Calculation of a discrimination index for each animal (**N**) demonstrated a preference for the displaced object above a chance level of 0.5. n= 10 (5 females, 5 males). **O-P**. C57BL/6J mice spent more time exploring the displaced (D) object compared to the non-displaced (ND) object during the test phase (**O**). Calculation of a discrimination index for each animal (**P**) uncovered a preference for the displaced object above chance levels (0.5). n=18 (9 females, 9 males). *p<0.05. Individual datapoints from female mice are depicted in orange and male mice in purple for transparency, but no sex differences were found.

The breadth and accessibility of genetic manipulations make the mouse an increasingly valuable model for the study of the biological basis of neurodevelopmental and neurological disorders. Although OiP recognition memory has been extensively studied in rats, few studies have examined it in mice. Importantly, when OiP memory is assessed in mice, it is either compounded with other types of recognition memory assessment within the same test session (Ricceri et al., 2000; Roullet et al., 2001; Calamandrei et al., 2002b, 2002a; Castilla-Ortega et al., 2012; Place et al., 2012; Sobin et al., 2017), or employ experimental designs not commonly found in the rat literature (Bonardi et al., 2016; Hall et al., 2016; Tsokas et al., 2016). Thus, a robust experimental design to exclusively assess OiP recognition memory in mice is currently missing, hindering the investigation of its underlying neural mechanisms.

Here, we first optimized the task design for the assessment of OiP recognition memory in the commonly used C57/129J and C57BL/6J mouse strains. We found that both strains performed optimally in a two-, but not four-object version of the OiP task. Using chemogenetic inhibition, we found that the iCA1, but not the mPFC or its input from iCA1, is necessary for two-object OiP recognition memory in mice. Our data identify a neural correlate for the expression of OiP memory in mice, expanding the available mouse behavioral testing repertoire and our understanding of the neural mechanisms underlying recognition memory processing.

## Methods

### Animals

Mice were either C57BL/6J (Jackson Laboratory) or a cross between C57BL/6J (maternal) x 129S1/SvImJ (paternal) strains (Jackson Laboratory; referred to as C57/129J for simplicity). Mice were bred at the University of Toronto Scarborough and kept on a 12h light/dark cycle (lights on at 07:00 h) with access to food and water *ad libitum*. Date of birth was designated postnatal day (P)0, with litter sizes ranging from 2 to 11 pups. At 21 days (P21), mice were weaned and housed in same-sex littermate groups of 2 to 5 mice. All experiments were conducted during the light cycle. Approximately equal numbers of females and males were used for all experiments. For all experiments, adult mice (P50-P80) were used. All animal procedures were approved by the Animal Care Committee at the University of Toronto.

### Apparatus & Objects

All recognition memory tests were conducted in a 30 x 30 x 30cm white plexiglass square chamber with a magnetic, glossy, removable base. The base consists of a 30 x 30cm plain white magnetic surface. Removable 3 x 3 cm squares were placed in the corner of the boxes to allow for accurate and consistent object placement and were removed prior to testing. Two of the four walls in the chamber contained visual cues that consisted of 8.5x11inch laminated white sheets with six large black dots (diameter 6.5 cm) at the north wall and 5 black stripes (17.5cm x 2cm) at the south wall, similar to Lesburguères et al., (2017). The chamber was elevated 41 cm off the floor with a camera mounted 75 cm above it using a wall mount rack. An external LED lamp (280 Lux) was located at the center of and 75cm above the four chambers to allow even distribution of light. The objects were designed using either SolidWorks ® (dome and step) or Tinkercad ® (pyramid and icosahedron) software and 3D printed using a LulzBot TAZ 6 3D printer with natural PLA or white PETG filament (see Inayat, Cruz-Sanchez et al. (2021) for freely available object designs). A round magnet (35mm diameter) was glued to the base of the objects to allow for stable attachment to the chamber floor. All objects had a pegged-surface with the following dimensions: 46 x 46 x 48mm (step), 47mm diameter x 48mm height (dome), 50.5 x 50.5 x 52mm (pyramid), and 50 x 50 x 50mm (icosahedron). Object designs were extensively piloted to generate objects that were (1) equal in surface area, (2) made of the same materials, and (3) for which the animals displayed no innate preference (Inayat, Cruz-Sanchez et al., 2021).

### Experimental Design and Statistical Analysis

#### Behavioral Testing

##### Handling & Habituation

Mice were handled and habituated to the behavioral chamber twice a day for four consecutive days prior to the day of testing for all experiments as in our previous work (Cruz-Sanchez et al., 2020). Handling took place in the testing room with a minimum 3h interval between handling sessions. Handling and habituation consisted of 5 minutes of handling followed by placement into the behavioral chamber for four minutes. A 4 x 4 cm weigh boat with 1-2g of rodent chow was placed at the center of the behavioral chamber during habituation to allow for better acclimation to the chamber. For identification purposes, mice were ear notched during surgery or were tail marked prior to handling.

##### General Procedures

To avoid confounds of repeated testing, dedicated cohorts of mice were used per experiment to ensure that each animal was only tested once. Behavioral chambers were cleaned with water between phases and subjects, and with 70% ethanol at the end of the day. For experiments that used a four-object paradigm, objects were placed in the northeast, northwest, southeast, and southwest corners of the behavioral chamber, 3cm away from each wall. The four objects were positioned in the same location during the sample phase for all animals (Lesburguères et al., 2017): the pyramid object at the northwest corner; the dome object at the northeast corner; the icosahedron object at the southeast corner; and the step object at the southwest corner (Figure 1A). Animals were placed into the chamber with their head facing either the north, east, west, or south wall of the behavioral chamber, in a balanced manner. To ensure an equal distribution of displaced object combinations as in previous OiP studies (Poucet et al., 1986; Granon et al., 1996; Dix and Aggleton, 1999; Barker and Warburton, 2015; Reichelt et al., 2015; Hall et al., 2016; Barker et al., 2017, 2020), displaced object pairs were counterbalanced across six possible combinations during the test phase (as suggested by Lesburguères et al. (2017): the pyramid and dome (top horizontal switch); the step and icosahedron (bottom horizontal switch); the dome and icosahedron (right vertical switch); the pyramid and step (left vertical switch); the dome and step (northeast-southwest diagonal switch); or the pyramid and icosahedron (northwest-southeast diagonal switch). For experiments that used a two-object paradigm, objects were placed in the northeast and northwest corners of the behavioral chamber, 3cm away from each wall. Displaced objects were counterbalanced for object type (i.e., dome vs step) and side (i.e., novelty on the right vs on the left).

##### Specific Procedures

##### Four-Object, One Sample Phase experimental design

This version of the object-in-place memory task comprised one sample phase followed by a test phase (Figure 1A). In the 5-minute sample phase, mice interacted with four different objects (pyramid, dome, step, and icosahedron), after which animals were removed and placed back into their home cage (Howland et al., 2012; Barker and Warburton, 2015). After a delay of 5 minutes (Barker et al., 2007; Barker and Warburton, 2008; Reichelt et al., 2015; Bonardi et al., 2021), mice underwent a 3-minute test phase (Dix and Aggleton, 1999; Barker et al., 2007, 2017; Barker and Warburton, 2008; Tsokas et al., 2016; Lesburguères et al., 2017) in which they were placed in the chamber with the same four objects, two of which had switched location. The displaced two-object combination was counterbalanced to ensure that all six possible combinations were utilized between animals (Lesburguères et al., 2017).

##### Four-Object, Two Sample Phase experimental design

This version of the OiP memory task comprised two sample phases followed by a test phase (Figure 1D). We followed the protocol described by Ricceri and colleagues, as most mouse studies that demonstrate successful expression of OiP memory with 3-5 objects do so using a minimum of 2 sample phases prior to testing (Ricceri et al., 2000; Calamandrei et al., 2002c, 2002a; Castilla-Ortega et al., 2012; Hall et al., 2016; Sobin et al., 2017). In each 2-minute sample phase, mice interacted with four different objects (pyramid, dome, step, and icosahedron) (Ricceri et al., 2000; Calamandrei et al., 2002c, 2002a). Once the mice completed both sample phases, they underwent a 3-minute test phase in which they were placed in the chamber with the same four objects, two of which had switched location. The interval between sample phases and between the sample and test phases was 2 minutes (Ricceri et al., 2000; Calamandrei et al., 2002c, 2002a), during which animals were placed back in their home cage. The displaced two-object combination was counterbalanced to ensure that all six possible combinations were used between animals (Lesburguères et al., 2017).

##### Two-Object experimental design

The two-object version of the object-in-place memory task comprised one sample phase followed by a test phase (Figure 1G). In the 5-minute sample phase (Davis et al., 2013), mice interacted with two objects (dome and step), after which animals were removed and placed back into their home cage. Object identity and location combinations were counterbalanced across animals. After a 5-minute delay (Davis et al., 2013; Bonardi et al., 2021), animals underwent a 3-minute test phase (Davis et al., 2013) in which they were placed in the chamber with two copies of the same object (two domes or two steps). Object type and novelty side (right vs left) at the test phase were also counterbalanced. This task was run in two versions: an egocentric or an allocentric version (Langston and Wood, 2010; Davis et al., 2013), which varied in the placement of the mouse at the start of the test phase to foster different spatial exploration strategies. In egocentric two-object experiments, animals were placed in the chamber with their head facing the wall located opposite the objects (i.e., south wall) for both the sample and test phases (Langston and Wood, 2010; Ainge et al., 2012; Davis et al., 2013). In experiments that used an allocentric two-object paradigm, animals were placed in the chamber with their head facing the south wall during the sample phase, and facing either the east or west wall (counterbalanced) during the test phase (Langston and Wood, 2010; Davis et al., 2013).

##### Behavioral analysis

Behavior was analyzed using ANY-maze® software (Stoelting). Exploratory activity was defined as in Leger et al., (Leger et al., 2013), similar to our previous work (Cruz-Sanchez et al., 2020). Briefly, object exploration was defined as an object-directed gaze while actively sniffing and/or pawing within 2 cm of the object. Sitting on top of the object while sniffing the surrounding air or chewing the object were not considered exploration. A discrimination index was calculated as a measure of relative novelty preference by dividing the amount of time spent exploring the displaced object(s) by the total time spent exploring both the displaced and non-displaced objects. Following Leger et al.’s (Leger et al., 2013) recommendation of using a 20 second criterion of exploration time for the test phase [adapted from (Arqué et al., 2008)], the analysis of the test phase of all experiments comprised the first 20 seconds of total interaction time with the objects, as in our previous work (Cruz-Sanchez et al., 2020). This is further supported by studies showing rodents demonstrate a higher preference for the novelty within the first 60 to 120 seconds of the test phase (Dix and Aggleton, 1999; Mumby, 2001; Mumby, Dave G., Stephanie, Gaskin., Glenn, Melissa J., Schramek, Tania E. & Lehmann, 2002; Cruz-Sanchez et al., 2020), which corresponds to when mice reached criterion in our sample. Importantly, we saw similar results when using the full 3 minutes for the four-object one-(unpaired t-test DI vs chance: *t*_(32)_=0.56, *p*=0.58), or two-sample phase (unpaired t-test DI vs chance: *t*_(38)_=0.86, *p*=0.39) and two-object (5min retention interval (RI) C57/129J unpaired t-test DI vs chance: *t*_(66)_=3.18, *p*=0.0022; 5min RI C57BL/6J unpaired t-test DI vs chance: *t*_(20)_=1.75, *p*=0.096; 1hr RI C57/129J unpaired t-test DI vs chance: *t*_(18)_=2.46, *p*=0.024; 1hr RI C57BL/6J unpaired t-test DI vs chance: *t*_(34)_=2.60, *p*=0.014) designs.

### Stereotaxic surgery and viral injections

Mice were anesthetized with isoflurane, and mounted onto a rodent stereotaxic apparatus (Stoelting) as in our previous work (Arruda-carvalho et al., 2017; Botterill et al., 2021). Approximately 0.2uL of pAAV5-hSyn-hM4D(Gi)-mCherry (≥ 7 x 10^12^vg/mL, Addgene # 50475), pAAV5-hSyn-mCherry (≥ 7 x 10^12^vg/mL, Addgene # 114472) or AAVrg-hSyn-Cre (≥ 1.8 x 10^13^ vg/ml, Addgene # 105553) virus was bilaterally infused onto the iCA1 (AP-2.75, ML-/+3.6, DV-3.00) or mPFC (spanning the infralimbic and prelimbic cortices; AP +2.08, ML +/-0.26, DV -2.7) using a 10ul NeurosSyringe (Hamilton Company) and motorized injector (Stoelting) as in our previous work (Arruda-Carvalho and Clem, 2014; Arruda-carvalho et al., 2017). Mice underwent stereotaxic surgery between P50-P70, with behavioral testing starting 10 days after surgery to allow for recovery. All mice that underwent surgery were given an intraperitoneal (i.p.) injection of the DREADD agonist compound 21 (C21, HelloBio #HB6124; 2mg/kg) 1-hour prior to the start of OiP testing.

### Slice Electrophysiology

We used slice electrophysiology to confirm the effect of C21 on hM4D-transfected neurons in the mPFC and iCA1 of adult C57BL/6J mice. Animals were anesthetized with isoflurane and perfused transcardially with ice-cold (0°C– 4°C) sucrose solution composed of (in mM): 197 sucrose, 11 glucose, 2.5 KCl, 1.25 NaH_2_PO_4_, 25 NaHCO_3_, 0.4 ascorbate, 1 CaCl_2_, 4 MgCl_2_ and 2 Na^+^ pyruvate. Acute 350mm coronal slices of mPFC or iCA1 were obtained using a VT1000S vibratome (Leica) and then incubated at 35°C for 25 min in the same solution, but with 50% standard artificial cerebral spinal fluid (ACSF) composed of (in mM): 120 NaCl, 2.5 KCl, 1.25 NaH_2_PO_4_, 25 NaHCO_3_, 11 glucose, 2 CaCl_2_, 1 MgCl_2_ and 2 Na^+^ pyruvate. Following recovery, slices were maintained at room temperature in standard ACSF.

Whole-cell current-clamp recordings were obtained from mCherry+ pyramidal neurons in layer 5 of the mPFC (prelimbic and infralimbic cortices) or mCherry+ pyramidal cells in the pyramidal layer of the iCA1 using borosilicate glass electrodes (3–5 MΩ) using an Olympus BX51WI microscope (Evident). Electrode internal solution was composed of (in mM): 127.5 K D-gluconate, 10 HEPES, 0.6 EGTA, 5 KCl, 2 MgCl_2_, 2 Mg-ATP, 0.3 Na_2_-GTP, 5 Na-phosphocreatine, and 0.4 Na-GTP. Data were low-pass filtered at 10 kHz and acquired at 10 kHz using Multiclamp 700B and pClamp 10 (Molecular Devices). Series and membrane resistance was continuously monitored, and recordings were discarded when these measurements changed by >20%. Baseline data was acquired by measuring resting membrane potential (RMP) and the number of action potentials evoked by a long (1s) depolarizing current pulse. Somatic current pulses were calibrated in the baseline period to trigger a minimum of 5 action potentials in hM4D-mCherry+ pyramidal neurons. After 10 minutes of stable neuronal discharge induced by a somatic depolarization, C21 (10 µM, HelloBio) (Jendryka et al., 2019) was bath applied for at least 10 minutes following which an additional recording of 10 minutes was compared to the baseline period.

### Statistical Analysis

Data are presented as mean ± SEM. All statistical analyses were performed in GraphPad Prism® version 9. Exploration time in the sample and test phases was analyzed using paired t-tests for non-surgery experiments, and with two-way, repeated-measures ANOVA followed by Sidak’s post-hoc tests in chemogenetic experiments. Within each treatment group, unpaired t-tests comparing DI to chance exploration level of 0.5 was calculated as in (Mumby, 2001; Gaskin et al., 2003; Bevins and Besheer, 2006). Object and side preference during the test phase (and sample phase for chemogenetic experiments) were assessed using a two-way, (repeated measures when necessary) ANOVA followed by Sidak’s post-hoc tests, and with a paired t-test for the sample phase in non-surgery groups (see detailed analysis below). For electrophysiology experiments, paired t-tests were performed to compare C21-mediated cell activity to baseline activity. Potential sex differences were first assessed using a two-way, (repeated measures when necessary) ANOVA and in the absence of effects, data were collapsed for subsequent analyses.

#### Detailed statistical analysis of potential side and object bias

We did not find innate side preferences (i.e. left vs right object, regardless of object type) during the sample phase [paired t-test; C57/129J 2-object: *t*_(24)_=0.12, *p*=0.90; C57BL/6J 2-object: *t*_(10)_=1.38, *p*=0.20; C57BL/6J 2-object 1hr retention interval (RI): *t*_(17)_=1.35, *p*=0.20; two-way ANOVA treatment x sample phase (left vs right exploration) interaction; iCA1 allocentric OiP inhibition: *F*_(1,19)_=0.98, *p*=0.34; mPFC allocentric OiP inhibition: *F*_(1,22)_=0.58, *p*=0.46; iCA1 egocentric OiP inhibition: *F*_(1,24)_=0.069, *p*=0.80; iCA1-mPFC allocentric OiP inhibition: *F*_(1,28)_=0.25, *p*=0.62], with the exception of the C57/129J two-object, 1hr RI group, which showed a bias toward the right object during the sample phase (paired t-test: *t*_(9)_=2.43, *p*=0.038). However, considering that this cohort demonstrated no innate object bias throughout the sample phase (Extended Data Figure 1-1J; paired t-test step vs dome: *t*_(9)_=1.45, *p*=0.18) or side preference within the chamber during the *test phase* (paired t-test (left vs right exploration): *t*_(9)_=1.50, *p*=0.17) and all objects were counterbalanced, it is unlikely this affected OiP memory expression in that cohort. During the test phase, we did not observe displaced side biases (i.e. preference for whether the displaced object was on the right vs left) in any of the experiments [two-way ANOVA displaced side x (displaced vs non displaced object exploration) interaction; C57/129J 2-object: *F*_(1,23)_=0.60, *p*= 0.45; C57BL/6J 2-object: *F*_(1,9)_=0.69, *p*=0.43; C57/129J 2-object 1hr RI: *F*_(1,32)_=0.66, *p*=0.42; C57BL/6J 2-object 1hr RI: *F*_(1,16)_=0.11, *p*=0.75; two-way ANOVA treatment DI x displaced side interaction; iCA1 allocentric OiP inhibition: *F*_(1,17)_ =3.29, *p*=0.087; mPFC allocentric OiP inhibition: *F*_(1,20)_ =2.21, *p*=0.15; iCA1 egocentric OiP inhibition: *F*_(1,22)_ =1.51, *p*=0.23; iCA1-mPFC allocentric OiP inhibition: *F*_(1,26)_ =1.05, *p*=0.31]. For allocentric experiments, we found no bias toward chamber placement (facing east or west wall) at the start of the test phase (two-way ANOVA treatment DI x chamber placement interaction; iCA1 allocentric OiP inhibition: *F*_(1,17)_ =4.01, *p*=0.061; mPFC allocentric OiP inhibition: *F*_(1,20)_ =0.061, *p*=0.11; iCA1-mPFC allocentric OiP inhibition: *F*_(1,26)_=0.13, *p*=0.72).

In addition to the sample phase object bias (e.g. dome vs steps) analyses featured in the results section, we also confirmed an absence of object type bias during the test phase across all experiments [two-way ANOVA object type x (displaced vs non displaced object exploration) interaction; 4-object: *F*_(5,11)_=2.14, *p*=0.14; 4-object, 2 sample phase: *F*_(5,14)_=0.41, *p*=0.84; C57/129J 2-object: *F*_(1,23)_=0.23, *p*= 0.60; C57BL/6J 2-object: *F*_(1,9)_=0.059, *p*=0.81; C57BL/6J 2- object 1hr RI: *F*_(1,16)_=0.45, *p*=0.51; two-way ANOVA treatment DI x object type interaction; iCA1 allocentric OiP inhibition: *F*_(1,17)_ =0.041, *p*=0.84; mPFC allocentric OiP inhibition: *F*_(1,20)_ =0.24, *p*=0.63; iCA1 egocentric OiP inhibition: *F*_(1,22)_ =1.24, *p*=0.28; iCA1-mPFC allocentric OiP inhibition: *F*_(1,26)_ =2.78, *p*=0.11], with the exception of the C57/129J two-object, 1hr RI group, which showed a preference toward the step object as the displaced object [two-way ANOVA object type (step vs dome) x (displaced vs non displaced object exploration) interaction: *F*_(1,8)-_

=19.04, *p*=0.0024). However, since objects were counterbalanced and there was no innate object preference during the sample phase (Extended Data Figure 1-1J; paired t-test step vs dome: *t*_(9)_=1.45, *p*=0.18), this should not affect our conclusions.

## Results

In all versions of the OiP task, mice undergo at least one sample or initial exposure phase, in which they are exposed to a set of objects. This is then followed by a test phase in which one or two of the objects has its identity/location switched, e.g. two out of four objects swap locations, such that their identity/location association is novel (Figure 1A). Increased exploration of the displaced object(s) is interpreted as evidence of OiP recognition memory, or the animal’s ability to successfully recall the association between the object’s identity and its prior location. To determine the neural correlates of OiP memory in mice, we first needed to establish a robust behavioral protocol to assess this type of recognition memory. Given our previous experience in running spontaneous recognition memory tasks using the C57/129J strain (Cruz-Sanchez et al., 2020; Inayat et al., 2021), we started by testing independent cohorts of C57/129J mice in the most common versions of the OiP recognition memory task (Ricceri et al., 1997, 2000; Dix and Aggleton, 1999; Calamandrei et al., 2002c; Barker and Warburton, 2008, 2015; Langston and Wood, 2010; Ainge et al., 2012; Davis et al., 2013; Ameen-Ali et al., 2015; Lesburguères et al., 2017; Sobin et al., 2017; Barker et al., 2017; Bonardi et al., 2021), spanning four (Dix and Aggleton, 1999; Barker and Warburton, 2008, 2015; Ameen-Ali et al., 2015; Benn et al., 2016; Barker et al., 2017; Lesburguères et al., 2017) to two objects (Langston and Wood, 2010; Langston et al., 2010b; Ainge et al., 2012; Davis et al., 2013; Ameen-Ali et al., 2015; Bonardi et al., 2021), as well as one to two sample phases (Ricceri et al., 1997, 2000; Calamandrei et al., 2002c, 2002a; Sobin et al., 2017).

### Four-object experimental design

We first tested C57/129J mice using the predominant OiP experimental design in rat studies, consisting of one sample phase and one test phase with four different objects (Barker et al., 2007; Barker and Warburton, 2008, 2011; Ameen-Ali et al., 2015; Barker and Warburton, 2015; Barker et al., 2017; Aggleton and Nelson, 2020; Barker et al., 2020). In this experimental design, mice undergo a sample phase in which they simultaneously explore four objects (dome, step, pyramid, icosahedron; Figure 1A) for five minutes. During the test phase, two of the objects switch locations (Figure 1A), and OiP recognition memory is measured by comparing the time mice spend exploring the displaced and non-displaced objects. Mice showed equivalent innate preference for all four objects in the sample phase, as measured by similar exploration times (Extended Data Figure 1-1A-B; one-way ANOVA, *F*_(3,48)_=2.44, *p*=0.076). In the test phase, C57/129J mice spent the same amount of time exploring the displaced and non-displaced objects (Figure 1B; paired t-test: *t*_(16)_=1.74, *p*=0.10). To determine whether mice were individually expressing a preference for the novel object-in-place contingency, we calculated a discrimination index by dividing the amount of time spent exploring the displaced objects by the total time spent exploring both the non-displaced and displaced objects, as in our previous work (Cruz-Sanchez et al., 2020). This analysis allows for the comparison of relative preference controlling for variability associated with individual differences in exploration. We found that the discrimination index was at chance level (0.5) (Figure 1C; unpaired t-test: *t*_(32)_=1.74, *p*=0.091), confirming a lack of preference for the displaced objects. We found no sex differences in OiP performance [two-way ANOVA sex x (displaced vs non displaced object exploration) interaction: *F*_(1,15)_=0.017, *p*=0.899]. These results show that C57/129J mice do not display OiP memory in a four-object, one sample phase experimental design.

While different rat strains perform well in the four-object, one sample phase design for OiP tests (Barker and Warburton, 2008, 2011; Howland et al., 2012; Barker and Warburton, 2015; Benn et al., 2016; Barker et al., 2017, 2020), mouse OiP studies using the CD1 strain have used a variation of that design in which animals are exposed to an additional, identical, sample phase (Figure 1D) (Ricceri et al., 2000; Calamandrei et al., 2002b, 2002a). To test whether additional exposure to the objects would facilitate OiP learning in C57/129J mice, we tested an independent cohort of mice using this experimental design (Ricceri et al., 2000; Calamandrei et al., 2002b, 2002a). C57/129J mice were exposed to four different objects during two identical, 2-minute sample phases with a two-minute interval (Figure 1D). Two minutes later, animals underwent a test phase in which two of the four objects switched locations. Similar to the original design, we found no innate preference for any of the sampled objects on the first (Extended Data Figure 1-1C-D; one-way ANOVA, *F*_(3,57)_=0.74, *p*=0.53) or second (Extended Data Figure 1-1E; one-way ANOVA, *F*_(3,57)_=0.67, *p*=0.57) sample phases. During the test phase, mice spent the same amount of time exploring the displaced and non-displaced objects (Figure 1E; paired t-test: *t*_(19)_=0.39, *p*=0.70). Similarly, the discrimination index did not differ from chance (Figure 1F; unpaired t-test: *t*_(38)_=0.31, *p*=0.76). We found no sex differences in discrimination using this version of the OiP memory task [two-way ANOVA sex x (displaced vs non displaced object exploration) interaction: *F*_(1,18)_ =0.99, *p*=0.33]. These results show that the additional sample phase was not sufficient to promote OiP recognition memory in C57/129J mice.

### Two-object experimental design

One main difference between the OiP task and other commonly used rodent recognition memory tasks is the number of objects used (Granon et al., 1996; Ricceri et al., 2000; Calamandrei et al., 2002c, 2002a; Langston and Wood, 2010; Ainge et al., 2012; Hall et al., 2016; Sobin et al., 2017; Bonardi et al., 2021). While frequently used spontaneous recognition tasks such as novel object recognition, object location, and temporal order recognition usually use two objects, OiP designs predominantly use four. Considering that the additional objects may increase task difficulty to a level that is not conducive with learning in this mouse strain, we next tested a two-object OiP design previously used in rats and two transgenic mouse studies (Eacott and Norman, 2004; Langston and Wood, 2010; Ainge and Langston, 2012; Davis et al., 2013; Bonardi et al., 2021). In this two-object experimental design, animals undergo one sample phase in which they are exposed to two different objects (dome and step) (Figure 1G). During the test phase, one object is replaced with an identical copy of the remaining, familiar object (two steps or two domes) (Figure 1G). Given that the two objects are identical during the test phase, OiP memory is interpreted as a preference for the replaced (referred to as displaced for consistency) object, i.e. the object whose identity-location association differs from the one present during the sample phase.

Mice showed no innate preference for object type during the sample phase (Extended Data Figure 1-1F-G; paired t-test: *t*_(24)_=0.06, *p*=0.96). During the test phase, mice spent more time exploring the displaced object (Figure 1H; paired t-test: *t*_(24)_=2.15, *p*=0.042). This preference was also expressed in the discrimination index (Figure 1I; unpaired t-test: *t*_(48)_=2.14, *p*=0.038). There were no sex differences in discrimination using this design [two-way ANOVA, sex x (displaced vs non displaced object exploration) interaction: *F*_(1,23)_ =0.37, *p*=0.55]. These results suggest that C57/129J mice successfully display OiP memory using the two-object experimental design. To further validate this OiP task design, we tested it using C57BL/6J mice, another commonly used mouse strain. We first confirmed that C57BL/6J mice did not show innate preference for object type during the sample phase (Extended Data Figure 1-1H; paired t-test: *t*_(10)_=0.1008, *p*=0.921). Similar to the C57/129J strain, C57BL/6J mice spent more time exploring the displaced object during the test phase (Figure 1J; paired t-test: *t*_(10)_=2.256, *p*=0.0477), demonstrating a preference for the displaced object (Figure 1K; unpaired t-test: *t*_(20)_=2.271, *p*=0.0344). These data indicate that the two-object OiP design produces robust OiP memory in two mouse strains.

Recognition memory studies show that different retention intervals may recruit different brain regions or neural circuits for a given task (Hammond et al., 2004; Norman and Eacott, 2004), including in the OiP task (Barker and Warburton, 2008, 2015, 2018; Barker et al., 2020). To test whether mice can retain two-object OiP memory for longer intervals, we next tested both C57/129J and C57BL/6J strains in this design with a 1-hour retention interval (Figure 1L). C57/129J mice showed an absence of object bias in the sample phase (Extended Data Figure 1-1I-J; paired t-test: *t*_(9)_=1.454, *p*=0.1799), and displayed increased exploration (Figure 1M; paired t-test: *t*_(9_)=2.689, *p*=0.0248) and preference (Figure 1N; unpaired t-test: *t*_(18)_=2.609, *p*=0.0178) for the displaced object. Similarly, C57BL/6J mice showed no bias during the sample phase (Extended Data Figure 1-1K; paired t-test: *t*_(17)_=1.843, *p*=0.0829), and successful OiP memory using the one-hour retention interval (Figure 1O; paired t-test: *t*_(17)_=2.761, *p*=0.0133; Figure 1P; unpaired t-test: *t*_(34)_=2..690, *p*=0.0110), indicating that both strains can retain two-object OiP memory for at least one hour.

### Chemogenetic inhibition of iCA1 impairs allocentric two-object OiP memory

While both the four- and two-object versions of the OiP tests are often treated as indistinctive assessments of OiP memory in the rodent literature, the question of whether they rely on the same brain regions has not been directly tested. Work by Langston and Wood confirmed that the hippocampus is necessary for two-object OiP memory in rats when the task is run in an allocentric configuration, in which the animals must rely on external cues to distinguish the objects (Langston and Wood, 2010). Furthermore, Barker and colleagues established that the four-object OiP task relies on the connection between iCA1 and the mPFC in rats (Barker et al., 2017). To test whether the two-object OiP task relies on these same brain regions in mice, we used designer receptors exclusively activated by designer drugs (DREADDs) to inhibit the iCA1, the mPFC and the iCA1-mPFC pathway in C57BL/6J mice.

Mice were first injected with AAV5-hM4D-mCherry or control AAV5-mCherry into the iCA1 (see Extended Data Figure 2-1 for viral targeting maps) and underwent the allocentric version of the two-object OiP task (Figure 2A; 5-minute retention interval). In this task design, the animals are placed in the chamber facing the wall opposite to the objects (i.e., south wall) in the sample phase, and placed facing the west or east wall of the arena in the test phase, precluding the use of an egocentric strategy to distinguish between the object placements (Langston and Wood, 2010; Davis et al., 2013). Mice were injected with the DREADD agonist C21 one hour prior to the start of behavioral testing (Figure 2A). Neither iCA1 mCherry or hM4D animals showed object bias in the sample phase (Figure 2B; two-way ANOVA, treatment x object type interaction; *F*_(1,19)_ =2.015, *p*=0.1719). During the test phase, while iCA1 mCherry animals spent more time exploring the displaced object, hM4D animals explored both objects equally (Figure 2C; two-way ANOVA main effect of object: *F*_(1,19)_ =8.221, *p*=0.0099, Sidak’s post-hoc test; mCherry: *t*_(19)_=2.681, *p*=0.0294, hM4D: *t*_(19)_=1.406, *p*=0.3209). Consistent with this, iCA1 hM4D animals failed to display a preference toward the displaced object as measured by their discrimination ratio (Figure 2D; unpaired t-test; mCherry: *t*_(20)_=3.370, *p*=0.003; hM4D: *t*_(18)_=1.182, *p*=0.2526). Importantly, iCA1 mCherry and hM4D animals did not differ in their total exploration time (unpaired t-test: *t*_(19)_=2.051, *p*=0.0543) or distance travelled (unpaired t-test: *t*_(19)_=0.3145, *p*=0.7566) during the test phase, suggesting that this effect was not driven by changes in motor behavior or motivation. These data show that chemogenetic inactivation of iCA1 impaired OiP recognition memory in the two-object version of this task.

**Figure 2.**
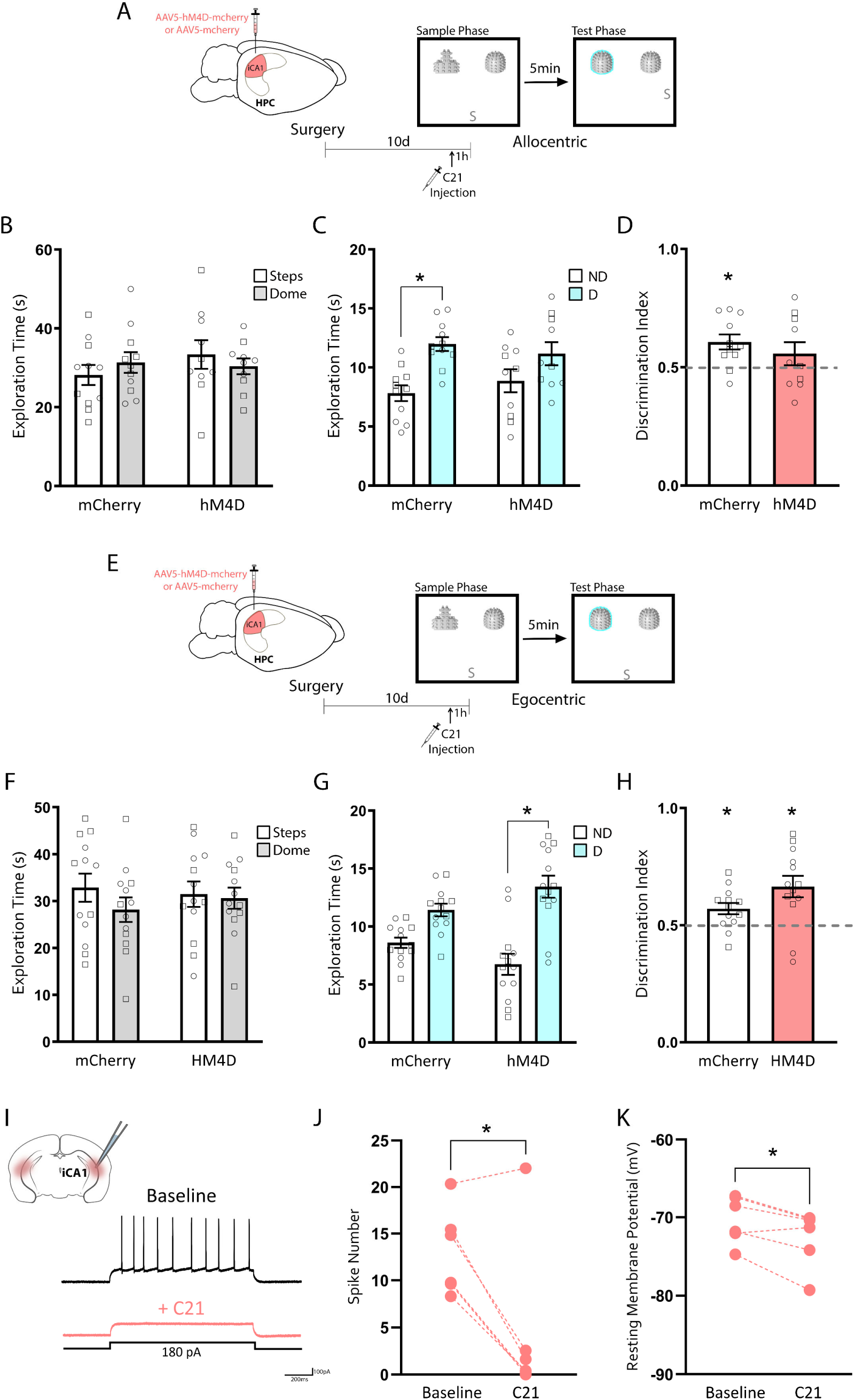
iCA1 activity is necessary for successful allocentric two-object OiP memory. **A-D**. iCA1 manipulation during allocentric two-object OiP. **A**. Experimental timeline. C57BL/6J mice were infused with AAV5-hM4D-mCherry or control AAV5-mCherry into the intermediate CA1 (iCA1) subdivision of the hippocampus (HPC). Ten days later, mice were injected with the DREADD agonist C21 one hour prior to testing in the allocentric version of the two-object OiP task. In the allocentric version of this task, animals were placed facing the south wall for the sample phase and facing the east or west wall for the test phase (depicted as S in the scheme). **B**. mCherry and hM4D iCA1 mice showed no object preference during the sample phase. **C**. During the test phase, mCherry iCA1 mice spent more time exploring the displaced (D) compared to the non-displaced (ND) objects, but hM4D mice displayed equal preference for both objects. **D**. Calculation of a discrimination index for each animal showed that hM4D iCA1 animals did not display a preference for the displaced object. mCherry: n=11 (5 females, 6 males); hM4D: n=10 (6 females, 4 males). **E-H**. iCA1 manipulation during egocentric two-object OiP. **E**. Schematic of the task design for the egocentric version of the two-object OiP task. Animals underwent identical procedures as in **A**, except that during the test phase they were placed in the chamber facing the same wall as the sample phase (depicted with an S in the scheme). **F**. mCherry and hM4D iCA1 mice showed no object preference during the sample phase. **G**. During the test phase, hM4D iCA1 egocentric mice spent more time exploring the displaced (D) compared to the non-displaced (ND) objects. **H**. Calculation of a discrimination index for each animal showed that both mCherry and hM4D iCA1 egocentric animals displayed a preference for the displaced object. mCherry: n=13 (6 females, 7 males); hM4D: n=13 (6 females, 7 males). Individual datapoints from female mice are depicted as circles and male mice as squares for transparency, but no sex differences were found. **I-K**. Electrophysiological validation of iCA1 DREADD experiments. **I**. Representative trace of action potentials recorded from an iCA1 hM4D-mCherry+ neuron at baseline and after application of C21**. J**. The number of action potentials evoked by somatic current injection (100pA) in iCA1 hM4D+ transfected neurons was significantly decreased following bath application of C21 (n=6 cells, 4 mice). **K**. The resting membrane potential (RMP) of iCA1 hM4D+ neurons became significantly more hyperpolarized following bath application of C21 (n=6 cells, 4 mice). *p<0.05.

Next, as a control, we ran mice in an egocentric version of the OiP task (Figure 2E-H), in which the animals are placed in the same location (facing the south wall) at the start of the sample and test phases. In that version of the OiP task, iCA1 inhibition did not affect preference for the displaced object, with iCA1 hM4D animals showing clear discrimination (Figure 2G; two-way ANOVA main effect of object: *F*_(1,24)_=20.63, *p*=0.0001, Sidak’s post hoc test; mCherry: *t*_(24)_=1.906, *p*=0.1327, hM4D: *t*_(24)_=4.518, *p*=0.0003; Figure 2H; unpaired t-test; mCherry: *t*_(24)_=2.921, *p*=0.0075, hM4D: *t*_(24)_=3.601, *p*=0.0014). Together, these data show that iCA1 activity is necessary for successful OiP recognition memory in the two-object version of the OiP task in mice when it is run in an allocentric configuration. We validated our chemogenetic iCA1 manipulation using whole-cell patch clamp recordings from hM4D+ neurons in the pyramidal layer of the iCA1 (Figure 2I-K). Bath application of C21 resulted in a decrease in spike number (Figure 2J; paired t-test: t_(5)_=3.854, p=0.012) and hyperpolarization of the resting membrane potential (Figure 2K; paired t-test: t_(5)_=3.245, p=0.0228) in iCA1 hM4D+ pyramidal neurons. These results confirm that acute application of C21 to hM4D-expressing iCA1 pyramidal neurons decreases their excitability.

### Chemogenetic inhibition of the iCA1-mPFC pathway or the mPFC does not affect allocentric two-object OiP memory

Given the involvement of the iCA1-mPFC pathway in four-object OiP memory in rats (Barker et al., 2017), we next tested whether this pathway would be necessary for two-object OiP recognition memory in mice using an intersectional approach. We injected a retrograde virus expressing Cre recombinase (AAVrg-Cre) in the mPFC (spanning the infralimbic and prelimbic cortices), and virus expressing Cre-dependent hM4D (AAV5-DIO-hM4D-mCherry) or mCherry (AAV5-DIO-mCherry) constructs in the iCA1 (Figure 3A-B), allowing for the inhibition of iCA1 neurons projecting to the mPFC. Mice were injected with C21 one hour prior to conducting the two-object, allocentric OiP task (Figure 3A). We found no object bias [Figure 3C; two-way ANOVA, significant treatment x (step vs dome exploration) interaction: *F*_(1,28)_ =0.5907, *p*=0.4486] in either control or hM4D mice during the sample phase. During the test phase, both hM4D and mCherry mice spent more time exploring the displaced object (Figure 3D; two-way ANOVA main effect of object: *F*_(1,28)_=14.0, *p*=0.0008, Sidak’s post hoc test; mCherry: *t*_(28)_=2.934, *p*=0.0132, hM4D: *t*_(28)_=2.377, *p*=0.0484). Both groups demonstrated a preference for the displaced object as measured by the discrimination index (Figure 3E; unpaired t-test; mCherry: *t*_(28)_=2.579, *p*=0.0155, hM4D: *t*_(28)_=2.743, *p*=0.0105). These data suggest that iCA1-mPFC activity is not necessary for the expression of two-object allocentric OiP memory in mice. Using whole-cell patch clamping to record from mCherry+ iCA1 pyramidal neurons of iCA1-mPFC hM4D mice (Figure 3F-H), we confirmed that C21 bath application led to a reduction in spike number (Figure 3G; paired t-test: t_(6)_=12.90, p<0.0001) and hyperpolarization (Figure 3H; paired t-test: t_(6)_=2.704, p=0.0354) of mPFC-projecting iCA1 pyramidal neurons expressing hM4D. While our data suggest that mice do not rely on the iCA1-mPFC pathway for successful OiP recognition memory, substantial research implicates the mPFC in OiP memory in rats (Barker and Warburton, 2008, 2009, 2011a, 2015; Savalli et al., 2015; Benn et al., 2016; Barker et al., 2017, 2020; Sabec et al., 2018), raising the question of whether the mPFC as a whole might still be involved in two-object OiP memory in mice. Therefore, we next infused the mPFC of mice with either control AAV5-mCherry or AAV5-hM4D-mCherry into the mPFC (see Extended Data Figure 4-1 for viral targeting maps), and tested them ten days later on the allocentric version of the two-object OiP task following C21 treatment (Figure 4A). Animals showed no bias for object type (Figure 4B; two-way ANOVA, treatment x object type interaction: *F*_(1,22)_ =0.888, *p*=0.3563) during the sample phase. During the test phase, both mPFC mCherry and hM4D animals showed a preference for the displaced object (Figure 4C; two-way ANOVA main effect of object: *F*_(1,22)_=26.93, *p*<0.0001, Sidak’s post hoc test; mCherry: *t*_(22)_=2.831, *p*=0.0194, hM4D: *t*_(22)_=4.447, *p*=0.0004; Figure 4D; unpaired t-test; mCherry: *t*_(24)_=2.589, *p*=0.0161; hM4D: *t*_(20)_=5.022, *p*<0.0001), indicating that mPFC inactivation does not affect OiP recognition memory in the two-object, allocentric version of the OiP. Validation experiments using patch clamp recordings showed that C21 triggered a reduction in spike number (Figure 4E-G; t_(8)_=3.157, p=0.0135) and hyperpolarization (Figure 4G; t_(8)_=2.771, p=0.0242) of mPFC pyramidal neurons expressing hM4D.

**Figure 3.**
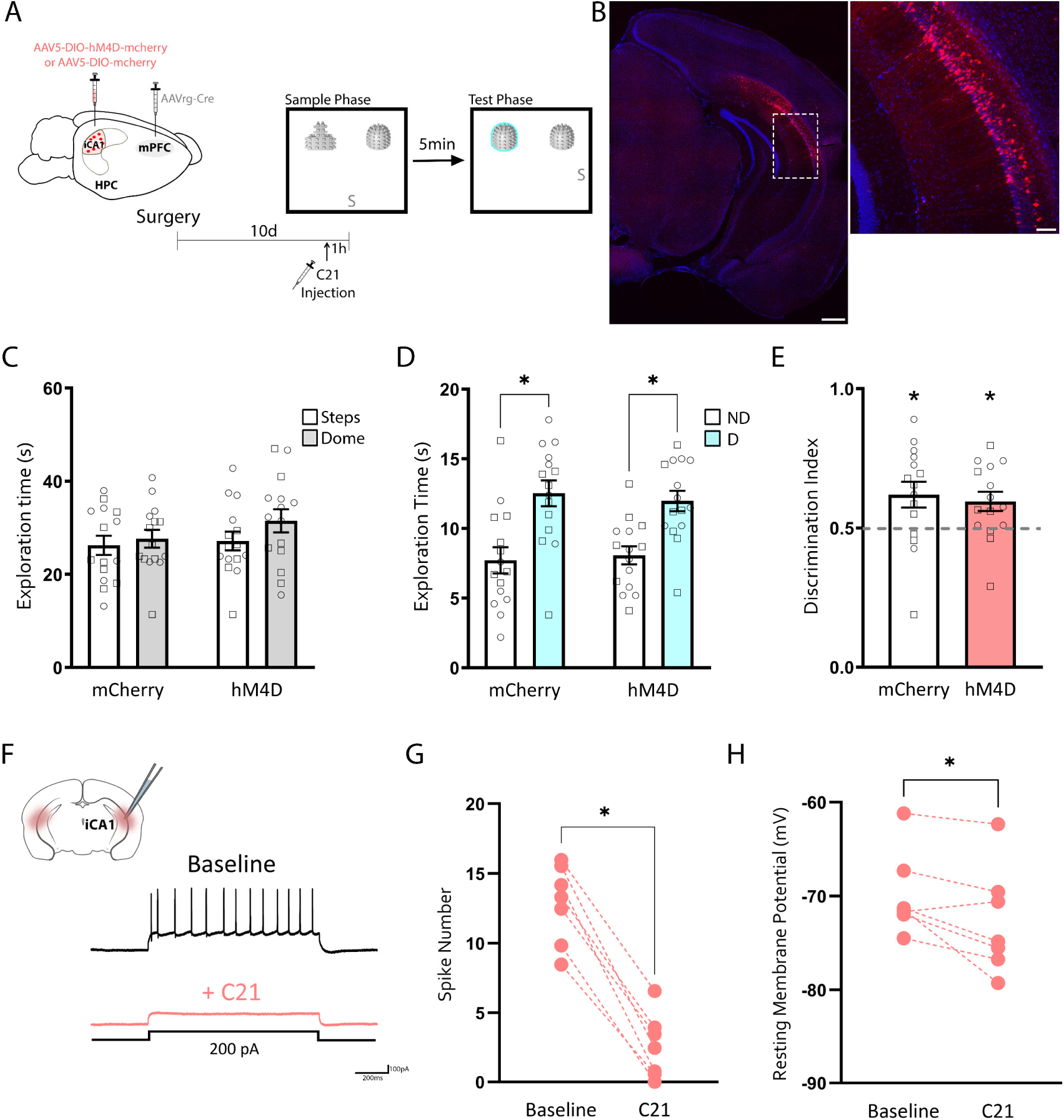
iCA1-mPFC activity is not necessary for successful two-object OiP memory at short intervals. **A**. Experimental timeline. C57BL/6J mice were infused with AAVrg-Cre into the medial prefrontal cortex (mPFC) and with either AAV5-DIO-hM4D-mCherry or control AAV5-DIO-mCherry into the intermediate CA1 (iCA1). Ten days later, mice were injected with the DREADD agonist C21 one hour prior to testing in the allocentric version of the two-object OiP task. **B**. Representative image showing transfected cells (red) in the iCA1, with insert magnified on the right image. Scale bars are 500mm (left) and 100mm (right). **C**. mCherry and hM4D iCA1-mPFC mice showed no object preference during the sample phase. **D**. During the test phase, both mCherry and hM4D iCA1-mPFC mice spent more time exploring the displaced (D) compared to the non-displaced (ND) objects. **E**. Calculation of a discrimination index for each animal showed that both mCherry and hM4D iCA1-mPFC animals displayed a preference for the displaced object. mCherry: n=13 (5 females, 8 males); hM4D: n = 11 (5 females, 6 males). Individual datapoints from female mice are depicted as circles and male mice as squares for transparency, but no sex differences were found. **F-H**. Electrophysiological validation of iCA1-mPFC DREADD experiments. **F**. Representative trace of action potentials recorded from a mPFC-projecting iCA1 hM4D-mCherry+ neuron at baseline and after application of C21. **G**. The number of action potentials evoked by somatic current injection (200pA) in hM4D-mcherry+ transfected neurons was significantly decreased following bath application of C21 (n=7 cells, 3 mice). **H**. The resting membrane potential (RMP) of DIO-hM4D+ neurons became significantly more hyperpolarized following bath application of C21 (n=7 cells, 3 mice). *p<0.05.

**Figure 4.**
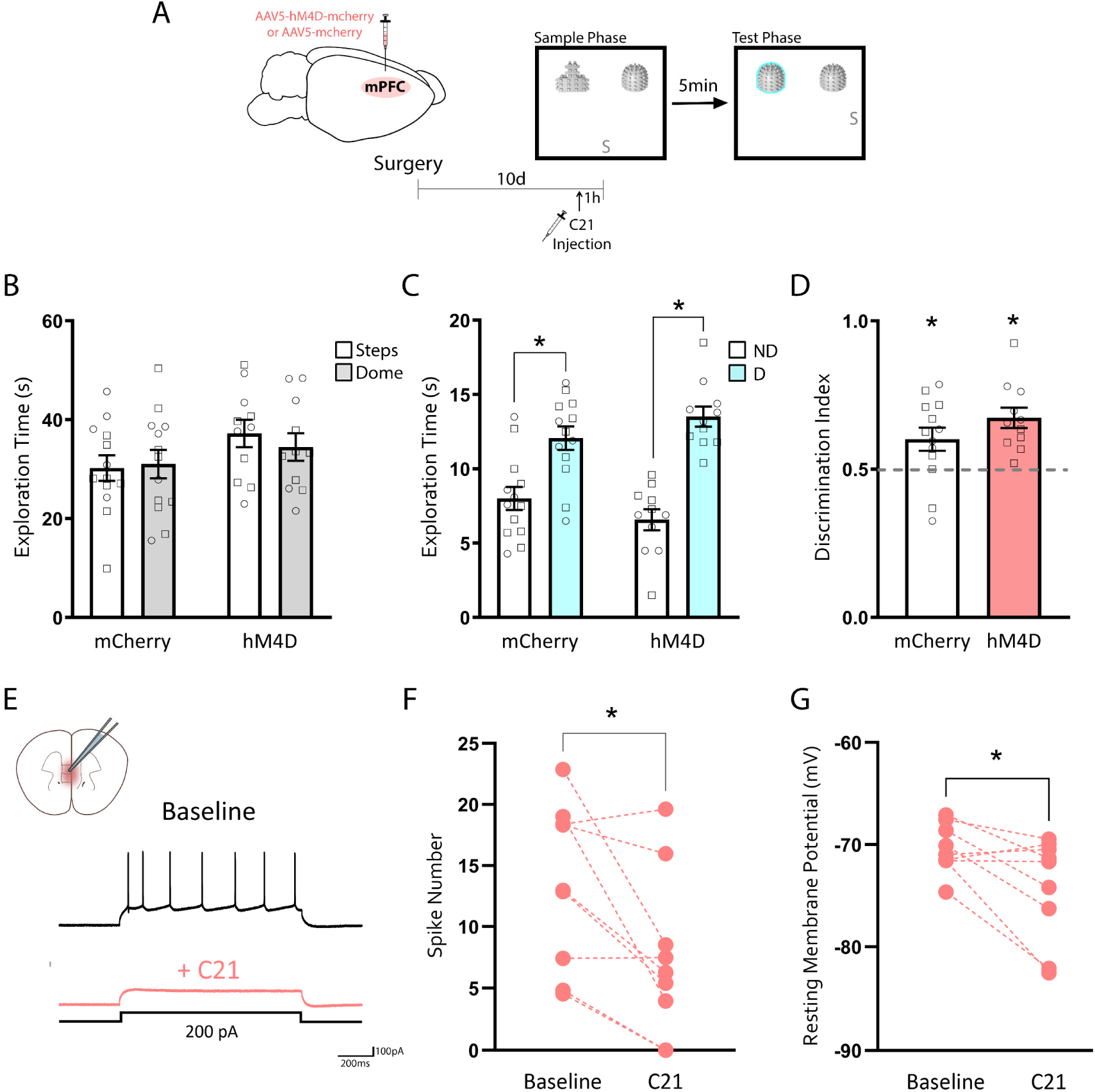
mPFC activity is not necessary for successful two-object OiP memory at short intervals. **A**. Experimental timeline. C57BL/6J mice were infused with AAV5-hM4D-mCherry or control AAV5-mCherry into the medial prefrontal cortex (mPFC). Ten days later, mice were injected with the DREADD agonist C21 one hour prior to testing in the allocentric version of the two-object OiP task. **B**. mCherry and hM4D mPFC mice showed no object preference during the sample phase. **C**. During the test phase, both mCherry and hM4D mPFC mice spent more time exploring the displaced (D) compared to the non-displaced (ND) objects. **D**. Calculation of a discrimination index for each animal showed that both mCherry and hM4D mPFC animals displayed a preference for the displaced object. mCherry: n=13 (5 females, 8 males); hM4D: n = 11 (5 females, 6 males). Individual datapoints from female mice are depicted as circles and male mice as squares for transparency, but no sex differences were found. **E-G**. Electrophysiological validation of mPFC DREADD experiments. **E**. Representative trace of action potentials recorded from a hM4D-mCherry+ neuron in the mPFC layer 5 at baseline and after application of C21. **F**. The number of action potentials evoked by somatic current injection (100pA) in hM4D+ transfected neurons was significantly decreased following bath application of C21 (n=9 cells, 6 mice). **G**. The resting membrane potential (RMP) of hM4D+ neurons became significantly more hyperpolarized following bath application of C21. (n=9 cells, 6 mice). *p<0.05.

## Discussion

To determine the neural correlates of OiP recognition memory in mice, we first established and validated a protocol to assess OiP recognition memory in C57/129J and C57BL/6J mouse strains across two retention intervals. We found that mice show OiP recognition memory in the two- but not four-object version of the OiP task, and that this memory is retained for at least one hour. Next, we chemogenetically inhibited the iCA1 of C57BL/6J mice during two-object OiP. We found that iCA1 activity is necessary for OiP recognition memory in the two-object allocentric version of this task with a short retention interval. In contrast, inhibition of iCA1 neurons projecting to the mPFC, or mPFC soma, did not affect OiP performance. Overall, our study is the first to establish a contribution of the iCA1 subregion to OiP memory in mice, expanding our understanding of the neural correlates of spatial recognition memory processing.

We found that C57/129J mice were unable to show OiP memory in the four-object version of the OiP task, even when exposed to an additional sample phase. While rats from different strains show robust four-object OiP memory across labs (Barker and Warburton, 2011a, 2015; Cost et al., 2012; Howland et al., 2012; Benn et al., 2016; Barker et al., 2017), few mouse studies showed successful four-object OiP memory, suggesting species differences in the ability to conduct this task (Ricceri et al., 2000; Calamandrei et al., 2002c, 2002a; Tsokas et al., 2016).

As the larger number of objects distinguishes the traditional OiP design from other commonly used two-object recognition tasks in mice, it is possible that the higher difficulty of the four-object OiP task may contribute to increased variability and sensitivity to subtle changes in experimental setup (Aggleton and Nelson, 2020). Interestingly, work by the Ricceri group found that CD1 mice were only able to display four-object OiP memory in late adulthood (P90) (Ricceri et al., 2000). Although the age of our adult animals varied between P50 and P80, we did not see any correlation between animal age and OiP performance in our data (age vs DI correlations p>0.2414), suggesting that strain differences between CD1 and C57/129J mice (Şik et al., 2003; van Goethem et al., 2012) might contribute to our negative findings. Importantly, the OiP task design used by the Ricceri group (Ricceri et al., 2000) and others (Sobin et al., 2017) comprised elements of both OiP and object location recognition memory. Specifically, out of the two displaced objects, one swapped position (as in OiP), while the other was moved to a new location previously unoccupied by an object (as in object location tasks). Since their analysis compounded both objects, we cannot exclude the possibility that discrimination in CD1 mice was facilitated or driven by the object location aspect of their task. To our knowledge, a single mouse study reported successful four-object OiP recognition memory without simultaneously testing for another type of recognition memory (Bonardi et al., 2016). In contrast to our experiments, this study used 21 week old mice, a longer sample phase and averaged object exploration across 2 repetitions of the OiP task, finding significant minute-by minute changes during the test phase (Bonardi et al., 2016). It is unclear whether the differences in task design or data analysis facilitated animal performance in that study.

We found that the iCA1 subregion of the hippocampus is necessary for two-object OiP in mice. Several studies implicate the hippocampus and its connections in OiP memory (Langston et al., 2010a; Barker and Warburton, 2011a; Barker et al., 2017). While Langston and Wood showed that whole hippocampal lesions impair two-object allocentric OiP performance in rats (Langston and Wood, 2010), our data restricted chemogenetic inhibition to the iCA1 subregion of the hippocampus, highlighting a potential functional specialization among hippocampal subregions in OiP as seen in rats (Barker et al., 2017). Notably, neither Barker and colleagues (Barker et al., 2017) or our study manipulated the ventral CA1, whose contribution to OiP memory is unclear. The intermediate hippocampus (iHPC) differs from its dorsal and ventral subdivisions in terms of cell population (Thompson et al., 2008; Cembrowski et al., 2016; Jin and Lee, 2021; Jarzebowski et al., 2022), efferent/afferent patterns (Fanselow and Dong, 2010) and synaptic activity (Izaki et al., 2001, 2002; Kawashima et al., 2006; Takita et al., 2010). Very few studies examined the functional role of the iHPC in memory and behavior. iHPC lesions are linked to deficits in one-trial spatial learning (Bast et al., 2009), and iCA1 activity is implicated in the encoding of reward-location associations and in correlating changes in motivational significance with spatial information (Jin and Lee, 2021; Jarzebowski et al., 2022). Together with our study, these data point to a unique role for the iHPC in the encoding of spatial location associations, particularly in the integration of spatial location with emotional and motivational information, assisting goal-directed navigation (Jin and Lee, 2021; Jarzebowski et al., 2022).

Our data showed a dissociation for the requirement of iCA1 between allocentric and egocentric configurations of the two-object OiP task. This is consistent with work in rats showing that the egocentric version of the two-object OiP task does not rely on the hippocampus (Langston and Wood, 2010). While the spatial complexity of four-object OiP clearly supports a need for the hippocampus in its completion (Langston et al., 2010a; Barker and Warburton, 2011a; Barker et al., 2017), these data in rats and mice reinforce the need for careful consideration of experimental parameters when running the two-object version of the OiP task.

We did not see a role for the iCA1-mPFC pathway in two-object OiP performance, in contrast to four-object OiP in rats (Barker et al., 2017). The iCA1 sends strong unidirectional projections to the prelimbic and infralimbic subdivisions of the mPFC (Takita et al., 2013), and the iHPC-mPFC pathway has been linked to working memory (Izaki et al., 2008) and valence changes in spatial navigation (Blanquat et al., 2013). Furthermore, OiP learning increases synchronization and theta phase locking between CA1 and mPFC in rats (Kim et al., 2011), with hippocampus-mPFC synaptic plasticity mechanisms being linked to OiP performance (Sabec et al., 2018), emphasizing a role for HPC-mPFC projections in OiP memory processing in rats. Perhaps even more surprisingly, mPFC inhibition also failed to affect two-object OiP performance in mice. Lesion, pharmacological and optogenetic mPFC loss- (Barker and Warburton, 2008; Cross et al., 2013; Savalli et al., 2015) and gain-of-function (Benn et al., 2016) manipulations in rats impair and facilitate four-object OiP memory at 5min retention intervals, respectively, pointing to a robust contribution of the mPFC to four-object OiP at short delays. Technical, species or experimental design (four- vs two-object) differences might be responsible for our negative results of iCA1-mPFC and mPFC inhibition in our task. For instance, studies suggest that mPFC activity is greater during certain spatial memory tasks when they are run in an egocentric configuration (De Bruin et al., 2001; Nieto-Escámez et al., 2002; Ma et al., 2003), raising the question of whether egocentric two-object OiP might require the mPFC. Ultimately, the two- and four-object versions of the OiP task may simply differ in their neural requirements, a possibility that remains to be directly tested within the same species.

Could another iCA1 downstream target mediate two-object OiP memory in mice? The perirhinal cortex (PRh) (Bussey et al., 2000; Barker and Warburton, 2008, 2009; Brown et al., 2012) and its connections with the HPC (Barker and Warburton, 2011) are necessary for four-object OiP expression in rodents, as well as OiP memory tasks using the Y maze in rats (Bussey et al., 2001) and OiP visual tasks in monkeys (Bachevalier and Nemanic, 2008). Interestingly, inhibition of the PRh during a two-object, radial maze OiP paired-associate task led to reduced dorsal CA1 activity (Lee and Park, 2013), suggesting that iCA1-PRh activity might be similarly affected in two-object OiP. However, the PRh only receives sparse projections from the iCA1 (Cenquizca and Swanson, 2007; Agster and Burwell, 2013), and a functional role for the iCA1-PRh pathway has not been described. In contrast, the iCA1 sends strong projections to the nucleus accumbens (NAc) (Kelley and Domesick, 1982; Groenewegen et al., 1999) and nucleus reuniens (Re) (McKenna and Vertes, 2004), two brains regions also implicated in OiP memory expression (Roullet et al., 2001; Barker and Warburton, 2018). Overall, while recent studies have made significant progress in elucidating specialized functions for the iCA1, the broader functional contribution of its downstream partners to behavior remains unknown.

Overall, this study identified a role for the iCA1 in two-object OiP recognition memory in mice, adding to a growing literature showing hippocampal subregion specialization in the requirement for the iCA1 in spatial associative memory. Our standardization of the OiP task with a two-object experimental design offers a robust, useful and versatile tool for cognitive assessment in mouse models of health and disease. Together, our findings expand the available mouse behavioral repertoire for cognitive behavioral testing and deepen our understanding of the neural basis of object identity-location associations in mice.

## Acknowledgements

This work was supported by a Natural Science and Engineering Council of Canada (NSERC) USRA for M.I, a CGS-D (CGSD3 - 534884 – 2019) for AC-S, as well as a grant from the Natural Science and Engineering Council of Canada (RGPIN-2017-06344) to MA-C. We acknowledge resources and support from the Centre for the Neurobiology of Stress (CNS) Core Facility at the University of Toronto Scarborough, and thank Durga Acharya and Bruno Chue for their help in the facility. A Canada Foundation for Innovation grant (#493864) was used to establish the Centre for the Neurobiology of Stress (CNS) Core Facility.

## Extended Data Figures and Legends

**Extended Data Figure 1-1.**
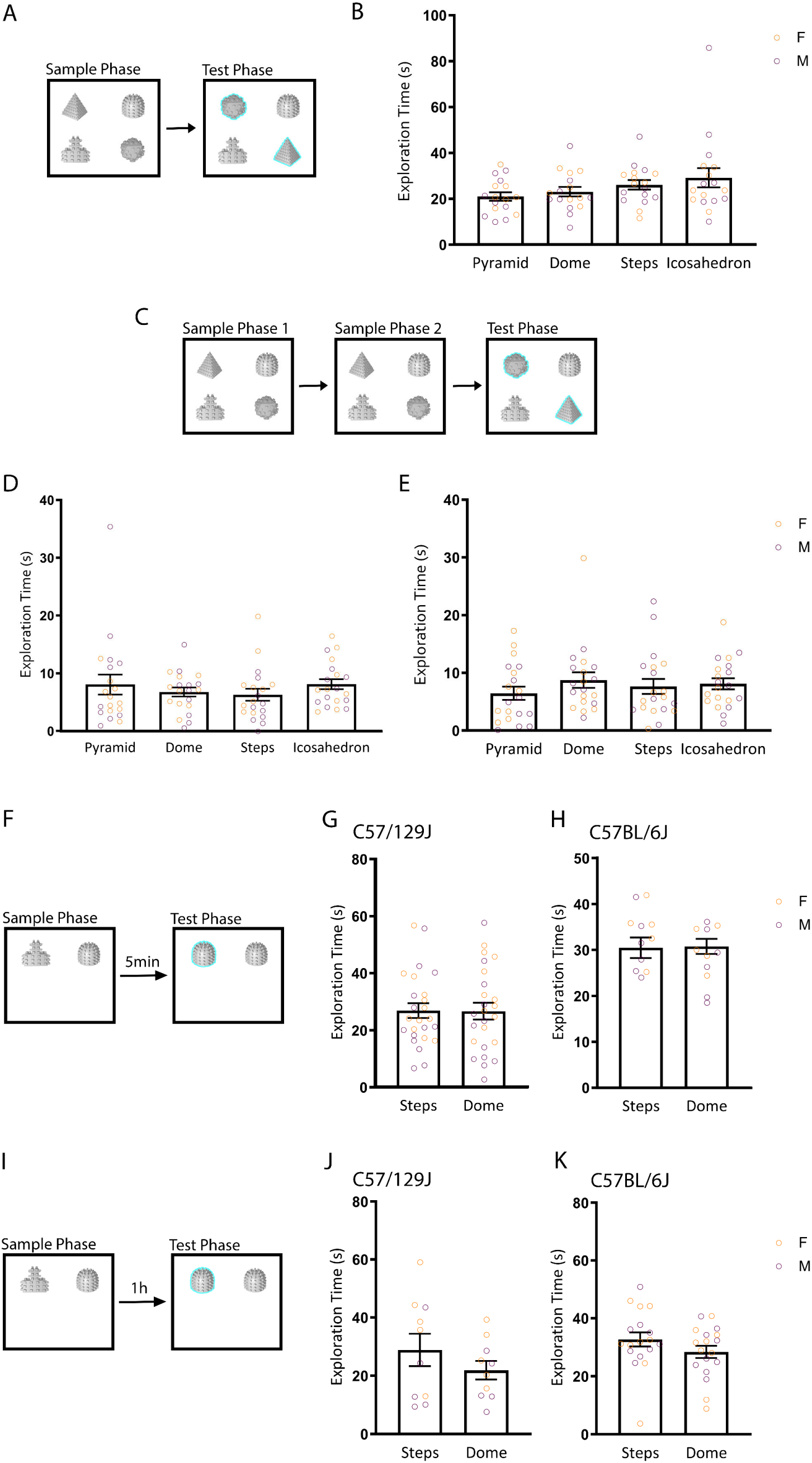
Sample phase object exploration across OiP task designs. **A-B**. Sample phase exploration time in the four-object, one sample phase OiP design. **C-E.** Sample phase exploration time in the four-object, two-sample phase OiP design. **D**. Exploration during sample phase 1. **E**. Exploration during sample phase 2. **F-H.** Sample phase exploration time in the two-object OiP design with a 5min retention interval. **G**. Sample phase exploration for C57/129J mice. **H**. Sample phase exploration for C57BL6/J mice. **I-K.** Sample phase exploration time in the two-object OiP design with 1h retention interval. **J**. Sample phase exploration for C57/129J mice. **K**. Sample phase exploration for C57BL6/J mice.

**Extended Data Figure 2-1.**
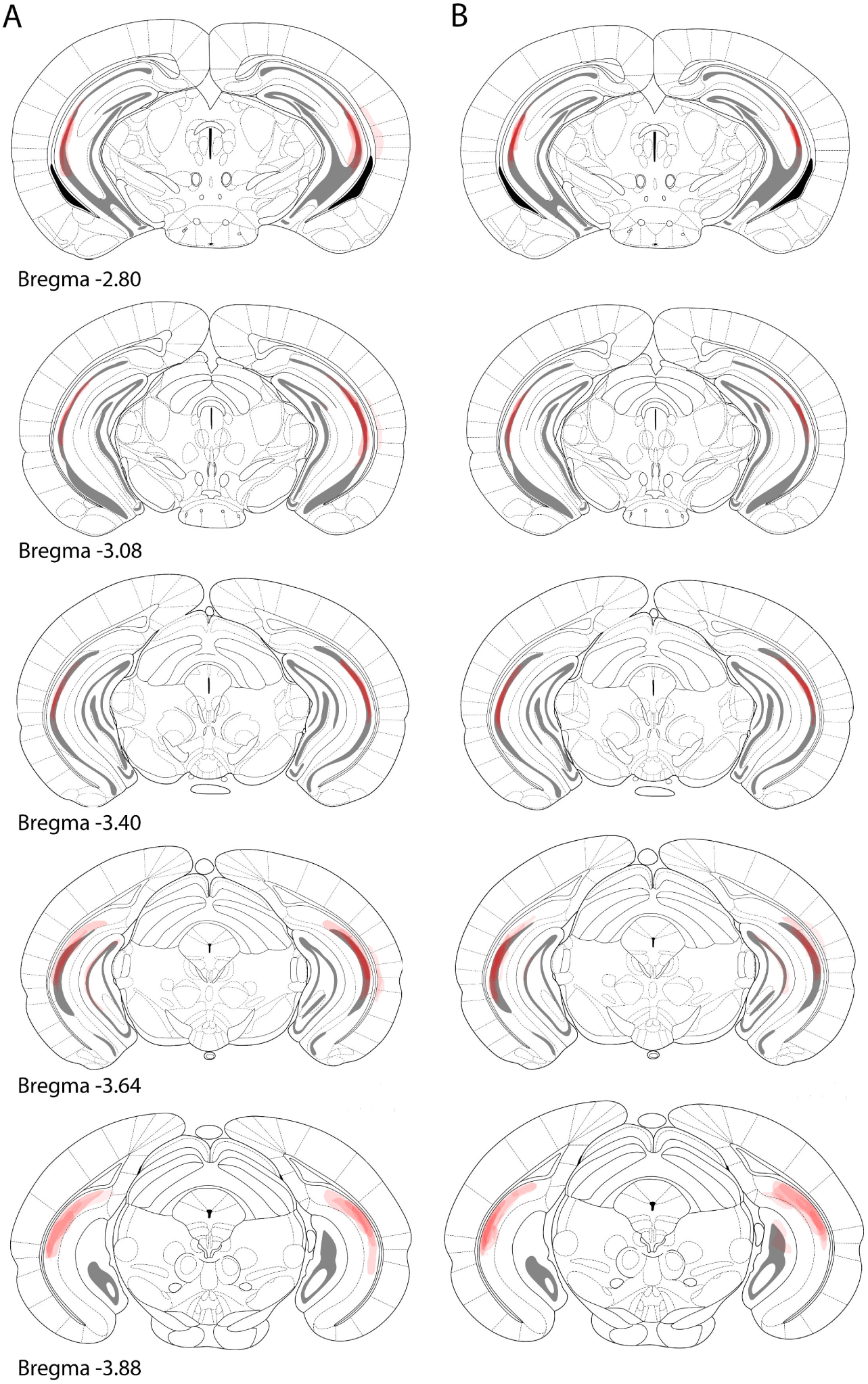
Viral expression maps of iCA1 manipulations. Diagrams depicting overlaid viral spread for all iCA1 hM4D mice that underwent the OiP task in the allocentric (**A**) or egocentric (**B**) configurations.

**Extended Data Figure 4-1.**
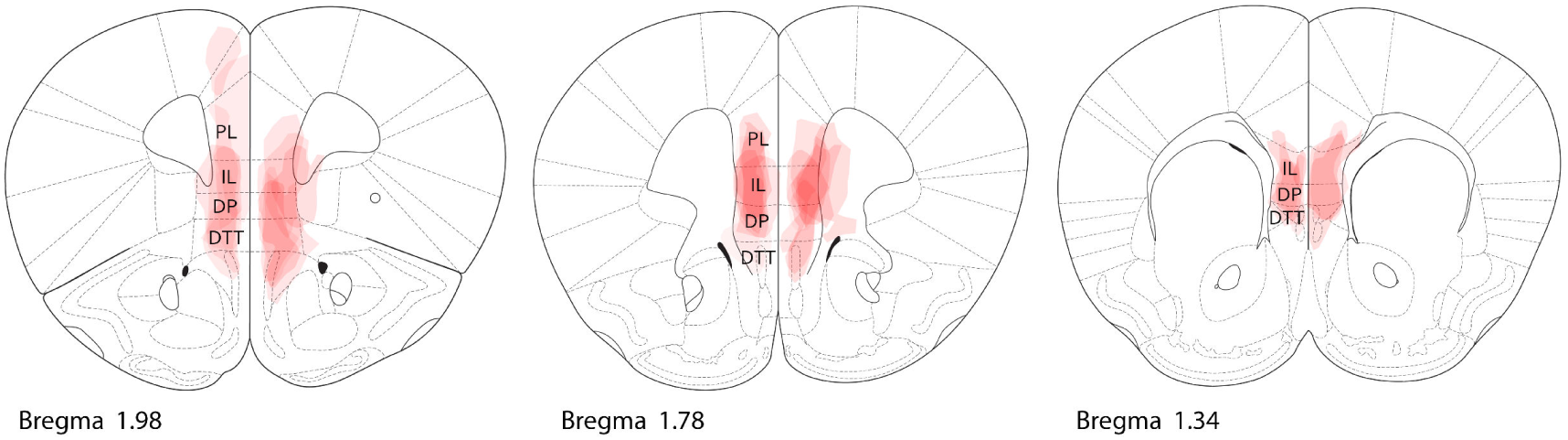
Viral expression maps of mPFC manipulations. Diagrams depicting overlaid viral spread for all mPFC hM4D infused animals. PL= Prelimbic cortex, IL=Infralimbic cortex, DP= Dorsal peduncular Cortex, DTT= Dorsal tenia tecta.

## References

Aggleton JP, Nelson AJD (2020) Distributed interactive brain circuits for object-in-place memory: A place for time? Brain Neurosci Adv 4:239821282093347.

Agster KL, Burwell RD (2013) Hippocampal and subicular efferents and afferents of the perirhinal, postrhinal, and entorhinal cortices of the rat. Behav Brain Res 254:50–64.

Ainge JA, Langston RF (2012) Ontogeny of neural circuits underlying spatial memory in the rat. Front Neural Circuits 6:1–10.

Ainge JA, Langston RF, Ascoli G, Mason G (2012) Ontogeny of neural circuits underlying spatial memory in the rat. Front Neural Circuits 6:1–10.

Ameen-Ali KE, Easton A, Eacott MJ (2015) Moving beyond standard procedures to assess spontaneous recognition memory. Neurosci Biobehav Rev 53:37–51.

Arqué G, Fotaki V, Fernández D, de Lagrán MM, Arbonés ML, Dierssen M (2008) Impaired spatial learning strategies and novel object recognition in mice haploinsufficient for the dual specificity tyrosine-regulated kinase-1A (Dyrk1A). PLoS One 3.

Arruda-Carvalho M, Clem RL (2014) Pathway-selective adjustment of prefrontal-amygdala transmission during fear encoding. J Neurosci 34:15601–15609.

Arruda-carvalho M, Wu W, Cummings KA, Clem RL (2017) Optogenetic Examination of Prefrontal-Amygdala Synaptic Development. J Neurosci 37:2976–2985.

Bachevalier J, Nemanic S (2008) Memory for Spatial Location and Object-Place Associations are Differently Processed by the Hippocampal Formation, Parahippocampal Areas TH/TF and Perirhinal Cortex. Hippocampus 18:64–80.

Ballendine SA, Greba Q, Dawicki W, Zhang X, Gordon JR, Howland JG (2015) Behavioral alterations in rat offspring following maternal immune activation and ELR-CXC chemokine receptor antagonism during pregnancy: Implications for neurodevelopmental psychiatric disorders. Prog Neuro-Psychopharmacology Biol Psychiatry 57:155–165.

Barker GR, Wong LF, Uney JB, Warburton EC (2020) CREB transcription in the medial prefrontal cortex regulates the formation of long-term associative recognition memory. Learn Mem 27:45–51.

Barker GRI, Banks PJ, Scott H, Ralph GS, Mitrophanous KA, Wong L, Bashir ZI, Uney JB, Warburton EC (2017a) Separate elements of episodic memory subserved by distinct hippocampal – prefrontal connections. Nat Neurosci 20.

Barker GRI, Banks PJ, Scott H, Ralph GS, Mitrophanous KA, Wong LF, Bashir ZI, Uney JB, Warburton EC (2017) Separate elements of episodic memory subserved by distinct hippocampal-prefrontal connections. Nat Neurosci 20:242–250.

Barker GRI, Bird F, Alexander V, Warburton EC (2007) Recognition memory for objects, place, and temporal order: a disconnection analysis of the role of the medial prefrontal cortex and perirhinal cortex. J Neurosci 27:2948–2957 Available at: http://www.ncbi.nlm.nih.gov/pubmed/17360918 [Accessed October 7, 2014].

Barker GRI, Warburton EC (2008) NMDA receptor plasticity in the perirhinal and prefrontal cortices is crucial for the acquisition of long-term object-in-place associative memory. J Neurosci 28:2837–2844.

Barker GRI, Warburton EC (2009) Critical role of the cholinergic system for object-in-place associative recognition memory. Learn Mem 16:8–11.

Barker GRI, Warburton EC (2011) When Is the Hippocampus Involved in Recognition Memory ? 31:10721–10731.

Barker GRI, Warburton EC (2015) Object-in-place associative recognition memory depends on glutamate receptor neurotransmission within two defined hippocampal-cortical circuits: A critical role for AMPA and NMDA receptors in the hippocampus, perirhinal, and prefrontal cortices. Cereb Cortex 25:472–481.

Barker GRI, Warburton EC (2018) A critical role for the nucleus reuniens in long-term, but not short-term associative recognition memory formation. J Neurosci 38:3208–3217.

Bast T, Wilson LA, Witter MP, Morris RGM (2009) From rapid place learning to behavioral performance: A key role for the intermediate hippocampus. PLoS Biol 7:0730–0746.

Baxter MG, Gaffan D, Kyriazis DA, Mitchell AS (2007) Orbital prefrontal cortex is required for object-in-place scene memory but not performance of a strategy implementation task. J Neurosci 27:11327–11333.

Benn A, Barker GRI, Stuart SA, Roloff EVL, Teschemacher AG, Clea Warburton E, Robinson ESJ (2016) Optogenetic stimulation of prefrontal glutamatergic neurons enhances recognition memory. J Neurosci 36:4930–4939.

Bevins RA, Besheer J (2006) Object recognition in rats and mice : a one-trial non-matching-to-sample learning task to study ‘ recognition memory .’ 1:1306–1311.

Blanquat PD Saint, Hok V, Save E, Poucet B, Chaillan FA (2013) Differential role of the dorsal hippocampus, ventro-intermediate hippocampus, and medial prefrontal cortex in updating the value of a spatial goal. Hippocampus 23:342–351.

Bonardi C, Pardon MC, Armstrong P (2016) Deficits in object-in-place but not relative recency performance in the APPswe/PS1dE9 mouse model of Alzheimer’s disease: Implications for object recognition. Behav Brain Res 313:71–81.

Bonardi C, Pardon MC, Armstrong P (2021) Time or place? Dissociation between object-in-place and relative recency in young APPswe/PS1dE9 mice. Behav Neurosci 135:39–50.

Botterill JJ, Khlaifia A, Walters BJ, Brimble MA, Scharfman HE, Arruda-Carvalho M (2021) Off-Target Expression of Cre-Dependent Adeno-Associated Viruses in Wild-Type C57BL/6J Mice. eNeuro 8:1–16.

Brown MW, Barker GRI, Aggleton JP, Warburton EC (2012) What pharmacological interventions indicate concerning the role of the perirhinal cortex in recognition memory. Neuropsychologia 50:3122–3140.

Burglen F, Marczewski P, Mitchell KJ, Linden M Van Der (2004) Impaired performance in a working memory binding task in patients with schizophrenia. 125:247–255.

Bussey TJ, Dias R, Amin E, Muir JL, Aggleton JP (2001) Perirhinal cortex and place-object conditional learning in the rat. Behav Neurosci 115:776–785.

Bussey TJ, Duck J, Muir JL, Aggleton JP (2000) Distinct patterns of behavioural impairments resulting from fornix transection or neurotoxic lesions of the perirhinal and postrhinal cortices in the rat. Behav Brain Res 111:187–202.

Calamandrei G, Rufini O, Valanzano A, Puopolo M (2002a) Long-term effects of developmental exposure to zidovudine on exploratory behavior and novelty discrimination in CD-1 mice. Neurotoxicol Teratol 24:529–540.

Calamandrei G, Valanzano A, Ricceri L (2002b) NGF induces appearance of adult-like response to spatial no v elty in 18-day male mice. 136.

Calamandrei G, Valanzano A, Ricceri L (2002c) NGF induces appearance of adult-like response to spatial novelty in 18-day male mice. Behav Brain Res 136:289–298.

Castilla-Ortega E, Pedraza C, Chun J, Fonseca FR de, Estivill-Torrús G, Santín LJ (2012) Hippocampal c-Fos activation in normal and LPA 1-null mice after two object recognition tasks with different memory demands. Behav Brain Res 232:400–405 Available at: 10.1016/j.bbr.2012.04.018.

Cembrowski MS, Bachman JL, Wang L, Sugino K, Shields BC, Spruston N (2016) Spatial Gene-Expression Gradients Underlie Prominent Heterogeneity of CA1 Pyramidal Neurons. Neuron 89:351–368.

Cenquizca L a, Swanson LW (2007) Spatial organization of direct hippocampal field CA1 axonal projections to the rest of the cerebral cortex. Brain Res Rev 56:1–26 Available at: http://www.pubmedcentral.nih.gov/articlerender.fcgi?artid=2171036&tool=pmcentrez&rendertype=abstract [Accessed May 24, 2013].

Connors EJ, Shaik AN, Migliore MM, Kentner AC (2014) Brain, Behavior, and Immunity Environmental enrichment mitigates the sex-specific effects of gestational inflammation on social engagement and the hypothalamic pituitary adrenal axis-feedback system. Brain Behav Immun 42:178–190.

Cost KT, Williams-Yee ZN, Fustok JN, Dohanich GP (2012) Sex differences in object-in-place memory of adult rats. Behav Neurosci 126:457–464.

Cross L, Brown MW, Aggleton JP, Warburton EC (2013) The medial dorsal thalamic nucleus and the medial prefrontal cortex of the rat function together to support associative recognition and recency but not item recognition. Learn Mem 20:41–50.

Cruz-Sanchez A, Dematagoda S, Ahmed R, Mohanathaas S, Odenwald N, Arruda-Carvalho M (2020) Developmental onset distinguishes three types of spontaneous recognition memory in mice. Sci Rep 10:1–11 Available at: 10.1038/s41598-020-67619-w.

Davis KE, Easton A, Eacott MJ, Gigg J (2013) Episodic-like memory for what-where-which occasion is selectively impaired in the 3xTgAD mouse model of Alzheimer’s disease. J Alzheimer’s Dis 33:681–698.

De Bruin JPC, Moita MP, De Brabander HM, Joosten RNJMA (2001) Place and response learning of rats in a Morris water maze: Differential effects of fimbria fornix and medial prefrontal cortex lesions. Neurobiol Learn Mem 75:164–178.

Dix SL, Aggleton JP (1999) Extending the spontaneous preference test of recognition: Evidence of object-location and object-context recognition. Behav Brain Res 99:191–200.

Eacott MJ, Norman G (2004) Integrated Memory for Object, Place, and Context in Rats: A Possible Model of Episodic-Like Memory? J Neurosci 24:1948–1953.

Fanselow MS, Dong HW (2010) Are the Dorsal and Ventral Hippocampus Functionally Distinct Structures? Neuron.

Gaffan D, Parker A (1996) Interaction of perirhinal cortex with the fornix-fimbria: Memory for objects and “object-in-place” memory. J Neurosci 16:5864–5869.

Gaskin S, Tremblay A, Mumby DG (2003) Retrograde and anterograde object recognition in rats with hippocampal lesions. Hippocampus 13:962–969.

Good MA, Barnes P, Staal V, McGregor A, Honey RC (2007) Context- but not familiarity-dependent forms of object recognition are impaired following excitotoxic hippocampal lesions in rats. Behav Neurosci 121:218–223.

Granon S, Save E, Buhot MC, Poucet B (1996) Effortful information processing in a spontaneous spatial situation by rats with medial prefrontal lesions. Behav Brain Res 78:147–154.

Griffiths S, Scott H, Glover C, Bienemann A, Ghorbel MT, Uney J, Brown MW, Warburton EC, Bashir ZI (2008) Expression of Long-Term Depression Underlies Visual Recognition Memory. Neuron 58:186–194.

Groenewegen HJ, Mulder AB, Beijer AVJ, Wright CI, Lopes Da Silva FH, Pennartz CMA (1999) Hippocampal and amygdaloid interactions in the nucleus accumbens. Psychobiology 27:149–164.

Hall JH, Wiseman FK, Fisher EMC, Tybulewicz VLJ, Harwood JL, Good MA (2016) Neurobiology of Learning and Memory Tc1 mouse model of trisomy-21 dissociates properties of short- and long-term recognition memory. Neurobiol Learn Mem 130:118– 128.

Hammond RS, Tull LE, Stackman RW (2004) On the delay-dependent involvement of the hippocampus in object recognition memory. Neurobiol Learn Mem 82:26–34.

Howland JG, Cazakoff BN, Zhang Y (2012) ALTERED OBJECT-IN-PLACE RECOGNITION MEMORY, PREPULSE INHIBITION, AND LOCOMOTOR ACTIVITY IN THE OFFSPRING OF RATS EXPOSED TO A VIRAL MIMETIC DURING PREGNANCY. NSC 201:184–198.

Inayat M, Cruz-Sanchez A, Thorpe HHA, Frie JA, Richards BA, Khokhar JY, Arruda-Carvalho M (2021) Promoting and optimizing the use of 3d-printed objects in spontaneous recognition memory tasks in rodents: A method for improving rigor and reproducibility. eNeuro 8.

Izaki Y, Takita M, Akema T (2008) Specific role of the posterior dorsal hippocampus-prefrontal cortex in short-term working memory. Eur J Neurosci 27:3029–3034.

Izaki Y, Takita M, Nomura M (2001) Mouse hippocampo-prefrontal paired-pulse facilitation and long-term potentiation in vivo. Neuroreport 12:1191–1193.

Izaki Y, Takita M, Nomura M (2002) Local properties of CA1 region in hippocampo-prefrontal synaptic plasticity in rats. Neuroreport 13:469–472.

Jarzebowski P, Hay YA, Grewe BF, Paulsen O (2022) Different encoding of reward location in dorsal and intermediate hippocampus. Curr Biol 32:834–841.e5.

Jendryka M, Palchaudhuri M, Ursu D, van der Veen B, Liss B, Kätzel D, Nissen W, Pekcec A (2019) Pharmacokinetic and pharmacodynamic actions of clozapine-N-oxide, clozapine, and compound 21 in DREADD-based chemogenetics in mice. Sci Rep 9:1–14.

Jin SW, Lee I (2021) Differential encoding of place value between the dorsal and intermediate hippocampus. Curr Biol 31:3053–3072.e5.

Kawashima H, Izaki Y, Grace AA, Takita M (2006) Cooperativity between hippocampal-prefrontal short-term plasticity through associative long-term potentiation. Brain Res 1109:37–44.

Kelley AE, Domesick VB (1982) The distribution of the projection from the hippocampal formation to the nucleus accumbens in the rat: An anterograde and retrograde-horseradish peroxidase study. Neuroscience 7:2321–2335.

Kentner AC, Khoury A, Lima Queiroz E, MacRae M (2016) Environmental enrichment rescues the effects of early life inflammation on markers of synaptic transmission and plasticity. Brain Behav Immun 57:151–160.

Kim J, Delcasso S, Lee I (2011) Neural correlates of object-in-place learning in hippocampus and prefrontal cortex. J Neurosci 31:16991–17006.

Langston R, Ainge J, Couey J, Canto C, Bjerknes T, Witter M, Moser E, Moser M (2010a) Development of the Spatial Representation System in the Rat. Science (80-) 328:1576–1581.

Langston RF, Stevenson CH, Wilson CL, Saunders I, Wood ER (2010b) The role of hippocampal subregions in memory for stimulus associations. Behav Brain Res 215:275– 291.

Langston RF, Wood ER (2010) Associative recognition and the hippocampus: Differential effects of hippocampal lesions on object-place, object-context and object-place-context memory. Hippocampus 20:1139–1153.

Lee I, Park SB (2013) Perirhinal cortical inactivation impairs object-in-place memory and disrupts task-dependent firing in hippocampal CA1, but not in CA3. Front Neural Circuits 7:1–10.

Leger M, Quiedeville A, Bouet V, Haelewyn B, Boulouard M, Schumann-bard P, Freret T (2013) Object recognition test in mice. Nat Protoc 8:2531–2537.

Lesburguères E, Tsokas P, Sacktor T, Fenton A (2017) The Object Context-place-location Paradigm for Testing Spatial Memory in Mice. Bio-Protocol 7:1–12.

Ma YY, Tian BP, Wilson FAW (2003) Dissociation of egocentric and allocentric spatial processing in prefrontal cortex. Neuroreport 14:1737–1741.

McKenna JT, Vertes RP (2004) Afferent projections to nucleus reuniens of the thalamus. J Comp Neurol 480:115–142.

Mitchnick KA, Creighton S, O’Hara M, Kalisch BE, Winters BD (2015) Differential contributions of de novo and maintenance DNA methyltransferases to object memory processing in the rat hippocampus and perirhinal cortex - a double dissociation. Eur J Neurosci 41:773–786.

Mitchnick KA, Creighton SD, Cloke JM, Wolter M, Zaika O, Christen B, Van Tiggelen M, Kalisch BE, Winters BD (2016) Dissociable roles for histone acetyltransferases p300 and PCAF in hippocampus and perirhinal cortex-mediated object memory. Genes, Brain Behav 15:542– 557.

Mitchnick KA, Mendell AL, Wideman CE, Jardine KH, Creighton SD, Muller AM, Choleris E, MacLusky NJ, Winters BD (2019) Dissociable involvement of estrogen receptors in perirhinal cortex-mediated object-place memory in male rats. Psychoneuroendocrinology 107:98–108.

Mumby, Dave G., Stephanie, Gaskin., Glenn, Melissa J., Schramek, Tania E. & Lehmann H (2002) Hippocampal Damage and Exploratory Preferences in Rats: Memory for Objects, Places, and Contexts. Learn Mem 9:49–57.

Mumby DG (2001) Perspectives on object-recognition memory following hippocampal damage: lessons from studies in rats. Behav Brain Res 127:159–181 Available at: http://www.ncbi.nlm.nih.gov/pubmed/11718890.

Nieto-Escámez FA, Sánchez-Santed F, De Bruin JPC (2002) Cholinergic receptor blockade in prefrontal cortex and lesions of the nucleus basalis: Implications for allocentric and egocentric spatial memory in rats. Behav Brain Res 134:93–112.

Norman G, Eacott MJ (2004) Impaired object recognition with increasing levels of feature ambiguity in rats with perirhinal cortex lesions. Behav Brain Res 148:79–91.

Place R, Lykken C, Beer Z, Suh J, McHugh TJ, Tonegawa S, Eichenbaum H, Sauvage MM (2012) NMDA signaling in CA1 mediates selectively the spatial component of episodic memory. Learn Mem 19:164–169.

Poucet B (1989) Object Exploration, Habituation, and Response to a Spatial Change in Rats Following Septal or Medial Frontal Cortical Damage. Behav Neurosci 103:1009–1016.

Poucet B, Chapuis N, Durup M, Thinus-Blanc C (1986) A study of exploratory behavior as an index of spatial knowledge in hamsters. Anim Learn Behav 14:93–100.

Reichelt AC, Killcross S, Hambly LD, Morris MJ, Westbrook RF (2015) Impact of adolescent sucrose access on cognitive control, recognition memory, and parvalbumin immunoreactivity. Learn Mem 22:215–224.

Ricceri L, Calamandrei G, Berger-sweeney J (1997) Different Effects of Postnatal Day 1 Versus 7 192 Immunoglobulin G-Saporin Lesions on Learning, Exploratory Behaviors, and Neurochemistry in Juvenile Rats. Ill:1292–1302.

Ricceri L, Colozza C, Calamandrei G (2000) Ontogeny of Spatial Discrimination in Mice : A Longitudinal Analysis in the Modified Open-Field with Objects. Dev Psychobiol 37:109– 118.

Roullet P, Sargolini F, Oliverio A, Mele A (2001) NMDA and AMPA antagonist infusions into the ventral striatum impair different steps of spatial information processing in a nonassociative task in mice. J Neurosci 21:2143–2149.

Sabec MH, Wonnacott S, Warburton EC, Bashir ZI (2018) Nicotinic Acetylcholine Receptors Control Encoding and Retrieval of Associative Recognition Memory through Plasticity in the Medial Prefrontal Cortex. Cell Rep 22:3409–3415.

Savalli G, Bashir ZI, Warburton EC (2015) Regionally selective requirement for D1/D5 dopaminergic neurotransmission in the medial prefrontal cortex in object-in-place associative recognition memory. Learn Mem 22:69–73.

Scott H, Smith A, Barker G, Uney J, Warburton E (2017) Contrasting roles for DNA methyltransferases and histone deacetylases in single-item and associative recognition memory. Neuroepigenetics.

Şik A, Van Nieuwehuyzen P, Prickaerts J, Blokland A (2003) Performance of different mouse strains in an object recognition task. Behav Brain Res 147:49–54.

Sobin C, Flores-Montoya MG, Alvarez JM (2017) Early chronic low-level Pb exposure alters global exploratory behaviors but does not impair spatial and object memory retrieval in an object-in-place task in pre-adolescent C57BL/6J mice. Neurotoxicol Teratol 61:104–114.

Takita M, Fujiwara SE, Izaki Y (2013) Functional structure of the intermediate and ventral hippocampo-prefrontal pathway in the prefrontal convergent system. J Physiol Paris 107:441–447.

Takita M, Izaki Y, Kuramochi M, Yokoi H, Ohtomi M (2010) Synaptic plasticity dynamics in the hippocampal-prefrontal pathway in vivo. Neuroreport 21:68–72.

Thompson CL, Pathak SD, Jeromin A, Ng LL, MacPherson CR, Mortrud MT, Cusick A, Riley ZL, Sunkin SM, Bernard A, Puchalski RB, Gage FH, Jones AR, Bajic VB, Hawrylycz MJ, Lein ES (2008) Genomic Anatomy of the Hippocampus. Neuron 60:1010–1021.

Tsokas P, Hsieh C, Yao Y, Lesburguères E, Wallace EJC, Tcherepanov A, Jothianandan D, Hartley BR, Pan L, Rivard B, Farese R V., Sajan MP, Bergold PJ, Hernández AI, Cottrell JE, Shouval HZ, Fenton AA, Sacktor TC (2016) Compensation for PKMζ in long-term potentiation and spatial long-term memory in mutant mice. Elife 5:1–22.

van Goethem NP, Rutten K, van der Staay FJ, Jans LAW, Akkerman S, Steinbusch HWM, Blokland A, van’t Klooster J, Prickaerts J (2012) Object recognition testing: Rodent species, strains, housing conditions, and estrous cycle. Behav Brain Res 232:323–334.

Wood SJ, Proffitt T, Mahony K, Smith DJ, Buchanan J-A, Brewer W, Stuart GW, Velakoulis D, McGorry P., Pantelis C (2002) Visuospatial memory and learning in first-episode schizophreniform psychosis and established schizophrenia : a functional correlate of hippocampal pathology ? Psychol Med 32:429–438.

Wout MV t., Aleman A, Kessels RPC, Kahn RS (2006) Object-location memory in schizophrenia: Interference of symbolic threatening content. Cogn Neuropsychiatry 11:272–284.

